# Microbial contaminants cataloged as novel human sequences in recent human pan-genomes

**DOI:** 10.1101/2020.03.16.994376

**Authors:** Mosè Manni, Evgeny Zdobnov

**Affiliations:** Department of Genetic Medicine and Development, University of Geneva Medical School, Geneva, Switzerland; Swiss Institute of Bioinformatics, Geneva, Switzerland

## Abstract

Human pan-genome studies offer the opportunity to identify human non-reference sequences (NRSs) which are, by definition, not represented in the reference human genome (GRCh38). NRSs serve as useful catalogues of genetic variation for population and disease studies and while the majority consists of repetitive elements, a substantial fraction is made of non-repetitive, non-reference (NRNR) sequences. The presence of non-human sequences in these catalogues can inflate the number of “novel” human sequences, overestimate the genetic differentiation among populations, and jeopardize subsequent analyses that rely on these resources. We uncovered almost 2,000 contaminant sequences of microbial origin in NRNR sequences from recent human pan-genome studies. The contaminant contigs (3,501,302 bp) harbour genes totalling 4,720 predicted proteins (>40 aa). The major sources of contamination are related to Rhyzobiales, Burkholderiales, Pseudomonadales and Lactobacillales, which may have been associated with the original samples or introduced later during sequencing experiments. We additionally observed that the majority of human novel protein-coding genes described in one of the studies entirely overlap repetitive regions and are likely to be false positive predictions. We report here the list of contaminant sequences in three recent human pan-genome catalogues and discuss strategies to increase decontamination efficacy for current and future pan-genome studies.

## Introduction

Hundreds of thousands of human genomes have been sequenced worldwide and ongoing consortiums and individual projects are sequencing many more^1,2^. The exponential growth of human whole-genome sequencing (WGS) data have already allowed the creation of population or cohort-specific human sequences to catalogue the genomic variation across different populations^3–5^. One of the main aims of pan-genome studies is to uncover novel non-reference sequences (NRSs) or variants not represented in the human reference genome (GRCh38). Although GRCh38 reference is of very high quality^6^, it contains gaps at repetitive and structurally diverse regions and it is likely biased towards individuals of European ancestry^7,8^. The use of NRSs from diverse ancestries can improve variant discovery and genotyping^7,9^, help identifying causal variants^10,11^, and ultimately facilitate disease research and clinical applications^12^. An early estimate quantified that the entire human pan-genome would contain an additional 19-40 Mb with respect to the reference sequence^13^. The largest pan-genome effort to date, consisting in sequencing 15,219 individuals from Iceland and focusing only on non-repetitive, non-reference (NRNR) sequences, uncovered a total of 326.6 Kbp in 3,719 NRNR sequences. More recent studies identified a larger amount of “novel” sequences. The African pan-genome^3^, compiled from 910 individuals, found ∼297 Mb of NRSs, while the analysis of 1,000 Swedish genomes^5^ uncovered 46 Mb of NRSs. The Han Chinese pan-genome found 29.5 Mb of NRSs and also identified 188 protein-coding genes^4^ defined as novel human genes. A vast fraction of NRSs in these catalogues corresponds to repetitive regions^11,14,15^, mostly centromeric, that are known to be underrepresented in the human reference genome^14^. A smaller fraction consists of NRNR sequences. Although a portion of NRSs may have no functional impact at the molecular or phenotypic level, some, especially NRNR sequences may represent causal variants underlying diseases^16,17^, representing a useful resource in genetic medicine. Human non-reference catalogues can also be highly valuable in metagenomics and microbiome research, as they can be used to aid the removal of human contaminant DNA from genomic and metagenomic datasets. Under such scenarios it is of paramount importance to efficiently remove, from raw datasets and assemblies, any off-target sequence which otherwise can be mistakenly annotated as human.

Identifying novel NRSs is of critical importance for understanding variation in human populations and we acknowledge the important contribution of such studies. Here we analyzed the sequences of the Pan-African pan-genome (APG)^3^, the Han Chinese pan-genome (HCP)^4^ and the Swedish NRSs^5^ and found evidence that a substantial fraction of NRNR sequences is of microbial origin. The presence of these sequences can yield misleading insights when treated and reported as human sequences, such as inflating the number of “novel” human sequences, overestimating the genetic differentiation among populations, and jeopardizing subsequent analyses that rely on these important resources. Despite each pan-genome study applied filters to remove contaminant sequences, it appears that the screening procedures adopted missed several contaminants. Here we provide a list of contaminant contigs that can be used to polish the respective NRSs catalogues and suggest a workflow that can be easily applied when checking for contamination in current and future pan-genome studies.

## Results

We analyzed the sequences of the Pan-African pan-genome (APG)^3^, the Han Chinese pan-genome (HCP)^4^ and the Swedish non-reference sequences (Swedish NRSs)^5^ (Supplementary Table 1). Our repeat annotation procedure masked 95% of APG, 87% of HCP and 65% of Swedish NRSs (Supplementary Table 2) confirming that a large fraction of these catalogs is mostly composed of low complexity and repetitive sequences. From a functional point of view, the remaining non-repetitive non-reference (NRNR) sequences represent a particularly interesting part of these collections. While we expected to encounter very little, if any, contamination, we identified 1,943 contaminants among these NRNR sequences, mostly in APG and HCP. We initially scanned the masked pan-genomes with Kaiju^18^ classifying sequences including bona fide human fragments (Supplementary Fig. 1). The Kaiju classified sequences were additionally compared to NCBI’s nt/nr databases of nucleotides and proteins, using BLASTN^19^ and DIAMOND^20^, respectively. Importantly, the non-repetitive nature of NRNR lowers the chances of getting spurious matches to non-human sequences. The results of BLASTn and DIAMOND searches were used to compute a Last Common Ancestor (LCA) for each sequence. For a true human sequence it would be unlikely to have a LCA in microbial taxa (e.g. all matches exclusively from bacterial species) unless the microbial sequences are the results of contamination from human DNA. Analyzing the hits and LCAs of each sequence we identified 1,475 contigs in APG, and 437 contigs in HCP which likely derived from microbial species (Fig. 1a,b,c; Supplementary Table 3; see paragraphs below and Methods section for details). Contaminant contigs are available in Supplementary File 1. These contigs harbour several putative protein-coding genes totalling 4,720 predicted proteins (>40 aa) of which 3,765 were either mapped to a bacterial orthologous group in OrthoDB v10 or have at least 1 hit to a Pfam domain (2,638 proteins; 56%) (Supplementary File 2; Supplementary Tables 3 and 4). Microbial sequences that matched the majority of contaminants were made publicly available in NCBI before 2018 (Fig. 1d). Accessions and publication dates of these sequences are reported in Supplementary Table 5. In Supplementary Table 3 we provide a list of contaminant sequences with corresponding LCA classifications and best hits from BLASTn and DIAMOND, and annotations from Pfam and InterPro databases. Supplementary File 3 illustrates the subset of contaminant contigs larger than 10 Kbp along with their annotated predicted proteins.

**Fig. 1:**
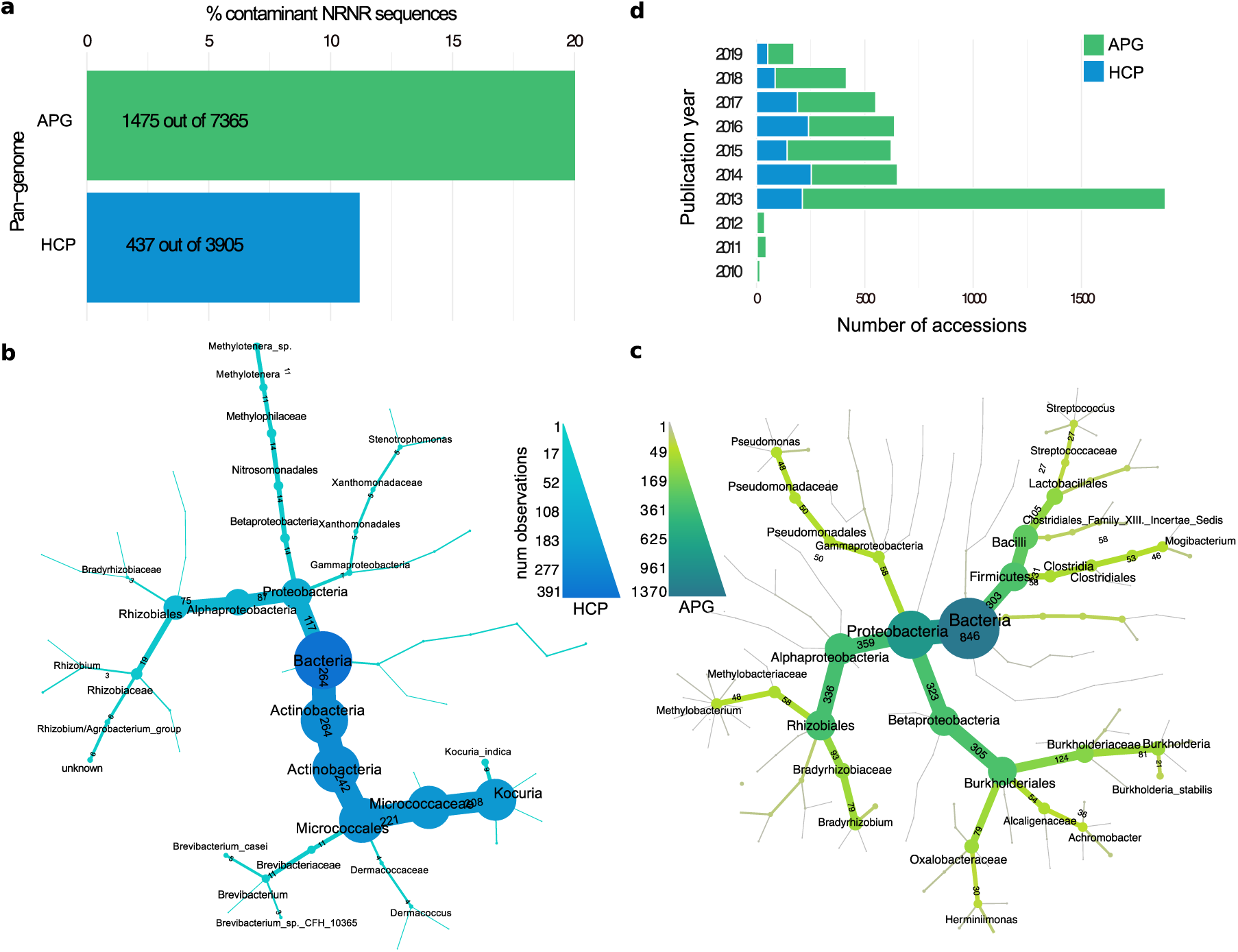
Contaminant sequences in non repetitive, non reference sequences of the APG and HCP. **a** Bar chart showing the percentage of contaminated contigs in NRNR sequences. The Swedish NRSs are not shown as only a small fraction of sequences were contaminants (see main text). **b-c** Taxonomic profiles of the last common ancestors of the contaminant contigs computed from DIAMOND results in APG and HCP respectively. Counts in panels (**b**) and (**c**) are based on the computed LCA of contigs that had at least 3 matches with >70% identity and >100 aa alignment length, thus excluding some of the contigs reported in panel (**a**). **d** Histogram displaying the publication date of publicly available microbial sequences in NCBI matching the contaminant contigs. Only the three oldest matches per sequence are plotted.

### Contamination in the African Pan-Genome (APG)

The African pan-genome was constructed from a dataset of 910 deeply sequenced individuals and uncovered 296 Mb of non-reference sequences^3^. According to our analyses the non repetitive fraction of the APG pan-genome consists of 7,365 contigs (5,85% of the total number) for ∼10.7 Mb (3.6% of the total size) of unmasked sequence. The rest of the pangenome consists of low-complexity regions or repetitive elements (Supplementary Table 2). Among the fraction of NRNR sequences we identified at least 1,475 (20%) contaminant contigs for a total length of 3,066,973 bp corresponding to 29% of the total size of the NRNR sequences (Fig. 1a,c). The contaminant contigs harbour several protein-coding genes totalling 4,011 predicted proteins (>40 aa) of which 2,350 (58%) gave hit to at least 1 Pfam domain. The longest contaminant contig (PDBU01047963.1) spans 61,265 bp and harbors 52 predicted proteins, of which 47 matched to known protein domains. The major source of contamination appears to be organisms from the order Rhyzobiales, Burkholderiales, and genuses *Phyllobacterium, Pseudomonas* and *Streptococcus* (Fig. 1c). Some contigs also resemble phage sequences (Supplementary Table 3). Fig. 2 highlights the pattern of contaminant sequences in the whole APG. Contaminants appear to cluster in blocks when all the contigs from the catalogue are displayed in numerical order (*i.e.* PDBU00000001.1, PDBU00000002.1, etc.). Contigs from individual “blocks” also appear to cluster by their taxon of origin, for example, the contaminant sequences in the range 45-50K mostly resemble Burkholderiales species (Supplementary Table 3). Given this pattern, it is likely that few individual human samples contributed to the contaminant contigs, which were grouped together when compiling the pan-genome.

**Fig. 2:**
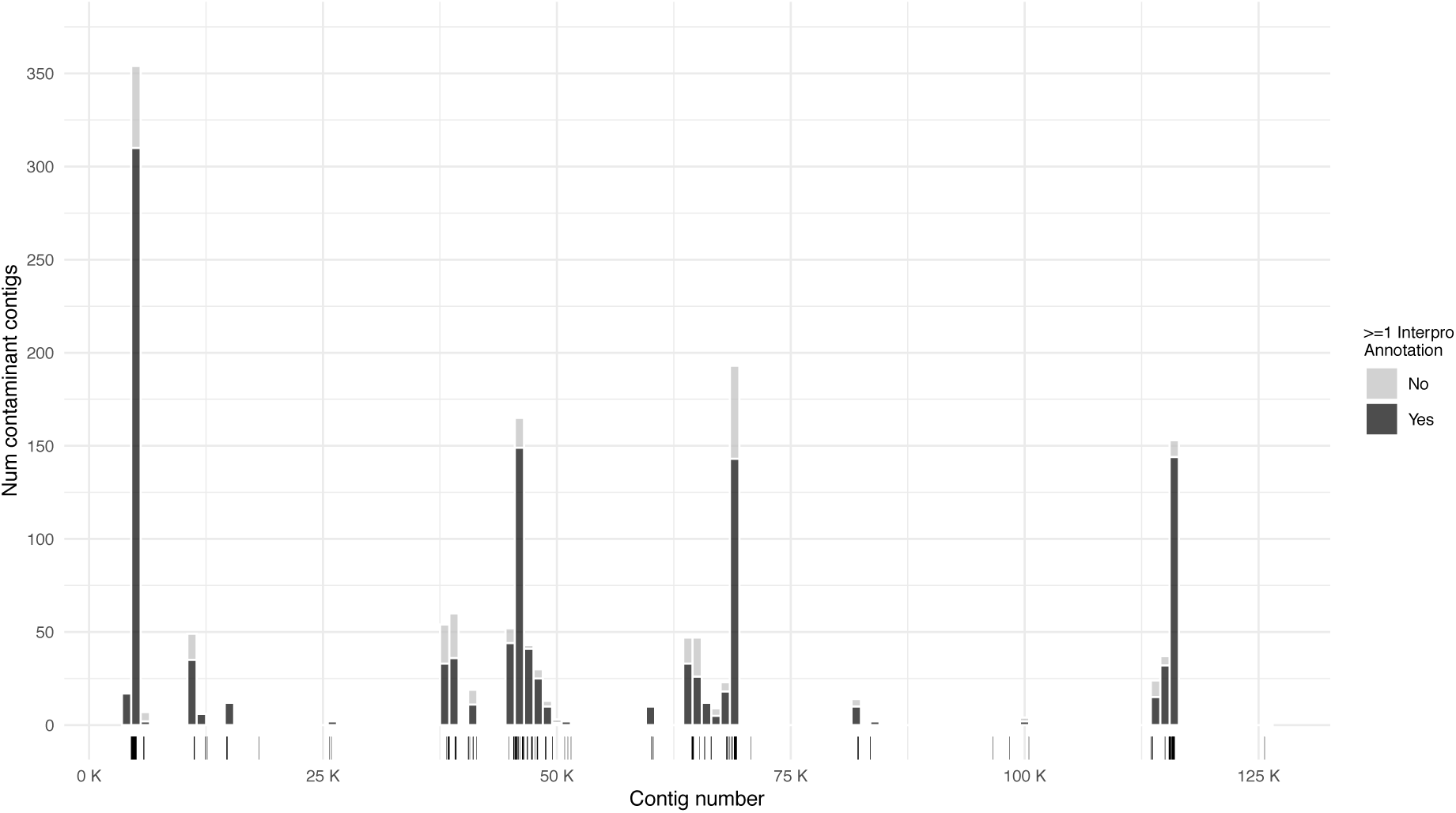
Histogram showing the occurrence of contaminant contigs in the APG. The x-axis corresponds to all the contigs (including the masked ones) reported in numerical order as they were numbered in the original study (*i.e.* PDBU00000001.1, PDBU00000002.1, etc.). On the y-axis is the number of contaminant sequences for bins of 1,000 contigs. The majority of contaminants appears to be grouped in 4 main clusters. These are likely to correspond to contigs deriving from individual human samples that carried the contaminants. Dark grey corresponds to the fraction of contigs harbouring at least 1 predicted protein with a match to a Pfam domain.

As example of contaminant contig, PDBU01060288.1 (25,547 bp) matches the genome of *Achromobacter xylosoxidans* strain FDAARGOS_147 in positions 3,319,249-3,344,796. This assembly is included in the Database for Reference Grade Microbial Sequences (FDA-ARGOS)^21^ which collects high quality bacterial genomes. While this assembly was deposited in NCBI in 2019, PDBU01060288.1 also gives hits to several Achromobacter and other bacterial genomes that were deposited in public databases since 2013. Under our settings, the PDBU01060288.1 LCA was defined at the *Achromobacter* genus level with 25 hits for the DIAMOND searches, and at the *Alcaligenaceae* family level with 2,954 hits from BLASTn results. The contig harbours 28 predicted protein-coding genes that have a match to known bacterial proteins (Fig. 3). These include PDBU01060288.1_6, a 441 amino acid (aa) long protein encoding for the TolB protein of the widely conserved Tol-Pal complex of Gram-negative bacteria; and PDBU01060288.1_14, a 704 aa long protein encoding for the prokaryotic elongation factor G involved in protein translation. Fig. 3 shows the alignment of the predicted TolB protein (PDBU01060288.1_6) from the contaminant contig with TolB proteins of other bacteria. Other examples of contaminant contigs are described in Supplementary Notes.

**Fig. 3:**
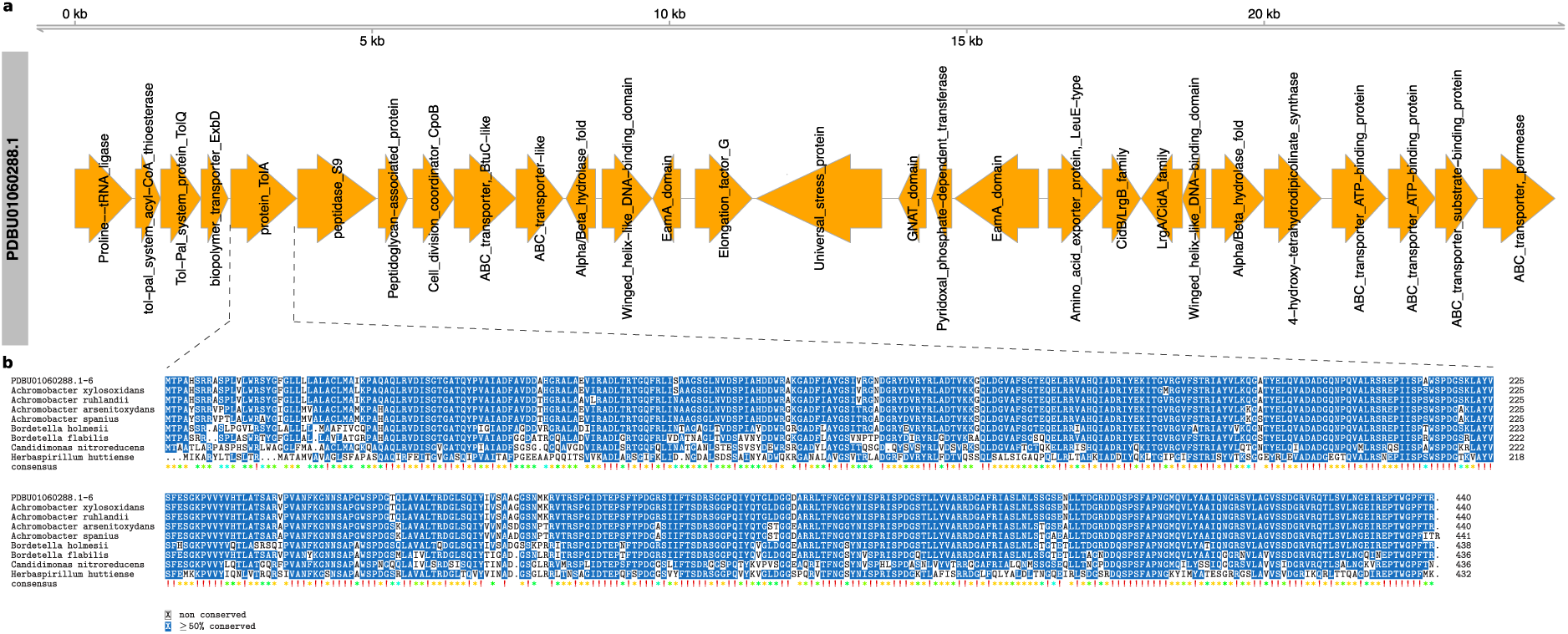
Example of contaminant contig PDBU01060288.1 in the APG. PDBU01060288.1 shares sequence homology with bacterial species of the *Achromobacter* genus. **a** The sequence map displays predicted protein-coding genes, in orange, along with their annotation derived from OrthoDB. **b** Multiple sequence alignment of the predicted protein PDBU01060288.1_6 with TolB proteins of different bacterial species.

### Contamination in the Han Chinese Pan-Genome (HCP)

The Han Chinese pan-genome was compiled from 275 individuals, including 185 newly sequenced^4^ and 90 previously published genomes^22^. The pan-genome was compiled using the HUman Pan-genome ANalysis (HUPAN) tool developed by the authors. Overall, the study identified 29.50 Mb of sequences missing from the GRCh38 (including patch sequences and alternative loci) and 188 novel human protein-coding genes. Approximately 88% of the 29.5 Mb was masked by our strategy. The unmasked fraction consists of 2.85 Mb (9.6% of the total size) in 3,905 contigs (13.8% of the total number of contigs). Of these NRNR sequences, at least 11.2% (437 sequences), corresponding to 405,766 pb (∼14.3% of the total size of NRNR sequences), were found to be contaminants (Fig. 1a,b). The major source of contamination (305 contigs) appears to be related to a species of *Kocuria* and derives from the 185 newly sequenced genomes. The other 132 contaminant contigs come from the 90 previously published genomes and appear to be related to various organisms mostly Rhizobiales species. Overall, the 437 contigs encode for 709 predicted proteins (>40 aa in length). Examples of contaminant contigs are described in Supplementary Notes.

From the HCP, the authors identified 188 novel protein-coding genes. 52 of these sequences derived from the 90 previously published genomes, and the rest from the 185 newly-sequenced genomes. Overall, the open reading frames of 176 of the novel protein-coding genes overlap repetitive or low-complexity regions (116,942 bp; 84.7%), including 152 that are entirely contained in such regions (Supplementary Table 6; Supplementary File 4). RepeatMasker classified 66,168 bp (47.95 %) (Supplementary Table 7) whereas dna-brnn, which specifically target satellites II/III (HSATII/III), identified 73,483 bp (53.3 %) (Supplementary Table 8). It is known that repetitive regions can result in false positive predictions of protein-coding genes^23,24^, and it appears this is the case for the majority of the 188 novel genes. These novel genes have been validated based on the evidence of transcription in human gastric tissues and other public RNA-seq datasets. However, also satellite repeats can also be transcribed^24,25^ and, in general, transcription per se is not indication of function at protein level. Of the 12 unmasked novel genes, 7 are of bacterial origin, specifically from the same taxa of Rhizobiales and Kokuria mentioned above. For example, Novel gene “185Genomes_NovelGene_004725” matches sequences annotated as molybdate ABC transporter permease subunit from several species of *Kocuria* (with identity ranging from 74 to 98%) with best match to sequences of *Kocuria indica* (98% identity; 110/112 aa). Matches for the other predicted genes are reported in Supplementary Table 9.

### Contamination in the Sweden NRSs

The Swedish NRSs catalogue was compiled from 1,000 samples and uncovered 46 Mb of novel sequences. We detected few contaminant contigs in this catalogue, specifically 31 contigs gave hit at the amino acid level (100% identity; >=99% query coverage) to sequences of streptomycin 3’’-adenylyltransferase (https://www.uniprot.org/uniprot/P0AG05), a protein that mediates bacterial antibiotic resistance to streptomycin and spectinomycin. This sequence has been used in numerous vector constructs. Indeed, at nucleotide level the contaminant contigs have 100% similarity over >=99% query coverage to suicide vectors pCD-RAsl1, pCD-RPyr4T, pCD-RPyr4, pCD-Dasl1. Fig. 4 shows the alignment of 13 Swedish NRSs to pCD-RAsl1 (KT176095.1). As we did not identify other contaminants from bacterial sequences it is likely that the source of contamination is indeed a vector that might have been introduced during the sequencing experiment or derived from cross-contamination.

**Fig. 4:**
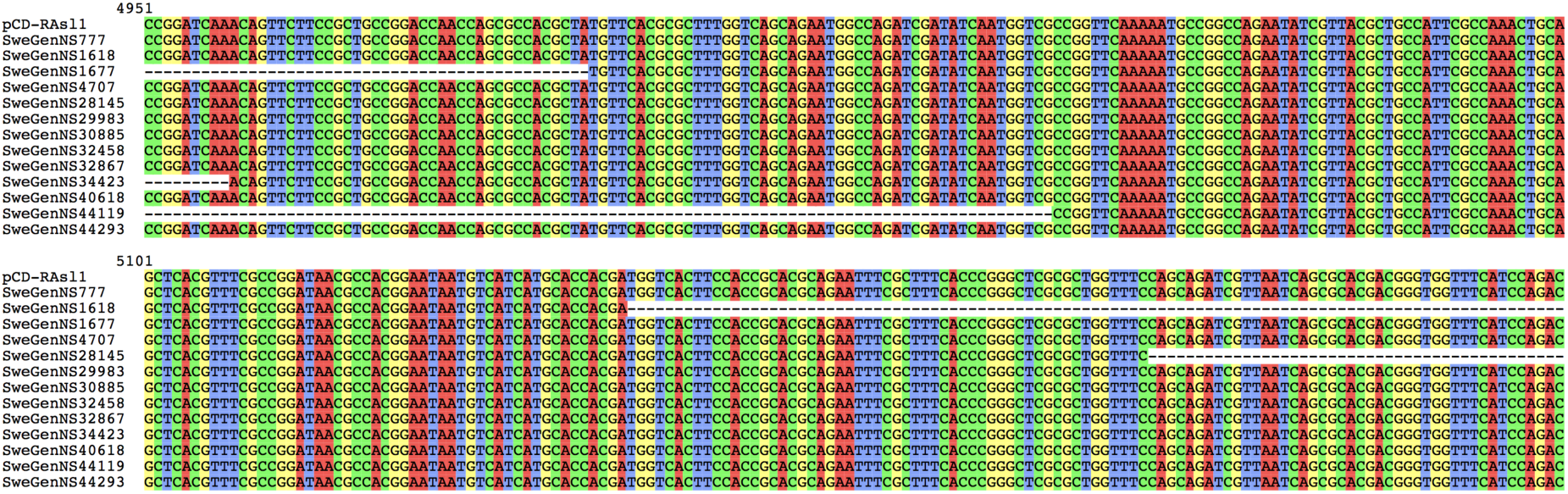
Multiple alignment showing the identical sequences of the Swedish NRSs and the suicide vector pCD-RAsl1. The homologous portion corresponds to the sequence of streptomycin 3’’-adenylyltransferase that is part of the vector construct.

### Human DNA in Microbial Genomes

During our analyses we additionally identified putative cases of contamination of human DNA in microbial sequences. Recently, Breitwieser et al.^25^ identified human DNA sequences in numerous microbial assemblies, that in some cases ended up being translated forming spurious bacterial proteins that have propagated in public databases^25^. The human reference genome (GRCh38) is commonly used for filtering human DNA from microbial genomic datasets. Reads corresponding to human sequences not represented in GRCh38 (such as the bona fide non-reference human sequences analyzed here) are not filtered and risk to be assembled and included in microbial assemblies. In our analyses such cases were easily identifiable because cross domain matches (e.g. to primates and bacteria) pushed the LCA level to “unknown” or “cellular organism”. In most of the cases, these sequences gave hits to spurious bacterial proteins from few species of bacteria when using DIAMOND searches (translated nucleotide versus database of proteins) while hit numerous primates DNA sequences with BLASTn searches. For example, 8 NRSs matched a *Butyrivibrio sp. XB500-5* protein annotated as homeobox domain-containing protein (RKM55982.1). We also observed 474 contigs with crossmatches between primates and sequences of *Plasmodium ovale wallikeri*. When using a protein database, these sequences matched three *Plasmodium* hypothetical proteins, SBT59247.1 (POVWA2_093830; 424/474 contigs), SBT58801.1 and SBT52172.1. However, these sequences also gave hit to numerous primates nucleotide sequences (Supplementary Table 10). This pattern suggests these are bona fide human sequences that were included in the Plasmodium assembly and ended up being translated into spurious proteins.

## Discussion

When human tissues are whole genome sequenced, any organism present in the sample will also be sequenced. High sequencing depths increase the chance of sequencing off-target DNA also from low microbial biomass samples such as blood which is commonly used as source of DNA for human WGS projects. Additional contaminants can derive from cross-contamination with other samples or sequencing runs, laboratory/ sequencing environments, reagents and nucleic acid extraction kits^30^. For a comprehensive review and recommendations on these issues see Eisenhofer et al.^30^. In a typical human WGS study the unmapped fraction of reads, that might also correspond to contaminants, is discharged. On the contrary, contamination can represent a major problem in both de novo and human pan-genome studies where the unmappable fraction of the data is kept.

We uncovered the presence of microbial DNA in recently compiled human pan-genomes by employing protein and nucleotide-based analyses following the workflow represented in fig. 5. Computation of LCAs allowed the identification of contaminant contigs and their taxonomic origin. As a result, the number of novel non-repetitive human sequences in these catalogues have been overestimated. The APG harbours the majority of the contaminants, while only few contaminant contigs were identified in the Sweden pan-genome study likely derived from vector constructs. Both were similarly compiled from ∼1,000 human samples. The Han Chinese pan-genome, compiled from 285 genomes, appeared to be mostly contaminated by sequences deriving from a single organism of the genus *Kokuria*.

**Fig. 5:**
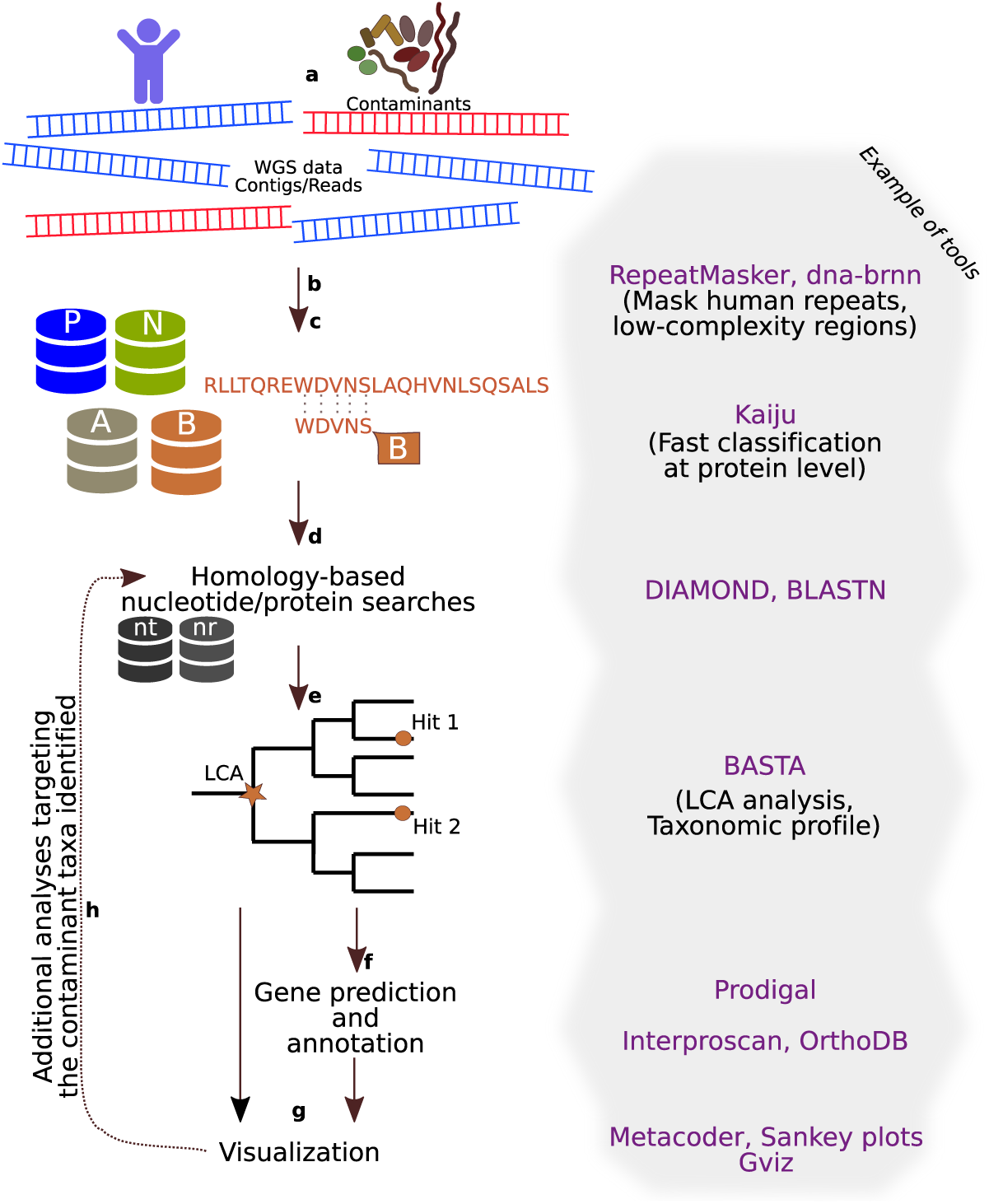
Example of workflow to detect contaminants in WGS data. **a** WGS data can contain contaminant sequences. **b** For human WGS, sequences can be optionally masked for human repeats and low-complexity regions, in order to reduce false positive matches like spurious hits to low-complexity regions or to human contaminant sequences present in microbial genomes (see main text for details). **c** Protein-based exact matches analyses coupled with the use comprehensive databases allow to classify sequences over longer evolutionary distances when compared to nucleotide-based analyses and can be used as first screening to detect potential contaminants. **d** Although too computationally demanding when done on the whole data set, homology-based analyses with large databases become feasible on the subset of sequences identified in the previous step. **e** Hits from the homology-based searches can be processed to compute the LCA and taxonomic profile of the contaminants. **f** Gene prediction and annotation of the contaminant sequences using protein/domain databases and mapping to orthologous groups. **g** Visualization of results to aid interpretation. For example, taxonomic profiles can be visualized with heat maps or Sankey plots, and contaminant sequences plotted along with their annotation and additional feature (e.g. GC%, read coverage). **h** Additional round of analyses can be carried out with an expanded set of sequences related to taxa identified as contaminants (e.g. with all available genomes) to further scan the input sequences.

The source material of the APG sequence data was blood^26^ and it is possible that microbial species were present in the original samples. This might be the case, among others, for *Burkholderia*, a diverse genus of gram-negative bacteria that can also infect humans (e.g. *Burkholderia stabilis)*, and members of the *Gemella* genus that are commensal organisms that occasionally cause blood infections^27–29^. The contaminant contigs related to *Kocuria* in HCP derived from samples of non-neoplastic gastric tissue. Speculating on the origin of these sequences, it is both possible that the *Kocuria* DNA was originally present in the samples or later introduced during WGS experiments through contaminating human skin, as a very closely related *Kocuria* strain has been isolated from the skin^31^. The other 90 previously published genomes from HCP were sequenced from cell lines DNA^22^, thus it is more likely the 132 contaminants derived from laboratory environments/kits or from cross contaminations. These and other contigs in APG appear to be related to taxa previously identified as common contaminants in sequencing experiments^30^, for instance, Bradyrhizobium and Methylobacterium.

The majority of the 188 genes in the HCP reported as novel protein-coding genes are unlikely to encode functional proteins as they appear to be the result of spurious gene predictions from repetitive/low complexity regions, and some turned out to be of bacterial origin.

The identification of non-reference human sequences is of prominent importance and will likely assume a central role in genetic medicine in the near future. Non-reference human sequences may represent causal variants underlying disease associations or may even harbor novel genes. For example, by analyzing WGS data of ∼15K Icelanders, Kehr et al.^17^ were able to identify 3,791 NRNR sequence variants, and demonstrated an association between a 766-bp NRNR (NRNR1361) and decreased risk of myocardial infarction.

To be a valuable resource in human genetics, human pan-genomes and catalogues of NRs should not contain contaminants, since overestimating the sequence and gene content diversity of human populations can result in misleading assumptions regarding the genetic background of specific populations and, in turn, bias genetic medicine perspectives. Pan-genomes are being used for studying population genetics and demographic histories of human populations, and the inclusion of non-human DNA can jeopardize population structure and differentiation estimates. Metagenomics analyses that rely on these human sequence catalogues to remove human host DNA from datasets can be affected by the presence of microbial contaminants as there is the risk of removing target microbial reads including potential pathogens. On the other hand, the availability and use of trusted comprehensive catalogues of human sequences can enable an efficient removal of human contaminant DNA from metagenomics datasets and alleviate contamination issues in microbial genomes.

The pan-genome studies considered here included a screening analysis for removing contaminant reads/ contigs. For example, Centrifuge^32^, a k-mer based tool commonly used for metagenomic classification, was used to scan reads in the APG study against the NCBI nucleotide database. However, the choice of the algorithm and database strongly influences the capacity to detect contaminant sequences. There are different reasons that may explain why the decontamination step missed the underlying reads/contigs of the contaminant taxa. One likely explanation is that the reference databases did not contain sequences of the contaminant organisms or they were too divergent to be classified by k-mer/nucleotide-based methods. Protein-based analyses such as Kaiju have the advantage of detecting divergent nucleotide sequences. The sequences are 6-frame translated and compared to protein databases allowing the detection of matches over longer evolutionary distances. This has a clear advantage over nucleotide-based classifiers that might fail to identify sequences if the database does not contain sequences from very closely related organisms (e.g. same species or strain) failing to detect contaminants from divergent taxa. The choice of databases is then critical regardless of the tool used for the classification, and despite many tools provide pre-built databases, these are often far from being comprehensive in terms of phylogenetic coverage, for example only including “complete genomes” from RefSeq.

Our observations suggest that removing contaminant DNA from human datasets still remains a challenging task. Based on our results, we recommend to employ both protein and nucleotide-based analyses coupled with the use of comprehensive databases to increase the detection of contaminants as illustrated in fig. 5. To make pan-genome data a trusted and useful resource for genetic medicine, in-depth pre and post-assembly analyses should be put in place to avoid contaminations in these important catalogues.

## Methods

We downloaded the African pan-genome (APG)^3^, Han Chinese pan-genome (HCP)^4^ and Swedish NRSs^5,33^ from the corresponding data sources (Supplementary Table 1). To focus on the non-repetitive fraction of these pan-genomes and to avoid spurious matches arising from low-complexity and repetitive regions we masked the pan-genomes with RepeatMasker v4.1.0 (http://www.repeatmasker.org) using the human repeats database (‘-species human’). To identify repeats and low complexity regions missed by RepeatMasker, we additionally run dna-brnn^15^, a program specific for human centromeric alpha satellite and satellite 2/3, dustmasker v1.0.0^34^ and bbmask (parameters: entropy=0.85 and sliding window=80bp) from BBTools version 38.71 (sourceforge.net/ projects/bbmap/). The coordinates corresponding to the masked sequences from each tool were combined and the pan-genomes masked with the maskfasta subprogram of bedtools v2.29.1^35^. We downloaded a local copy of the nt and nr databases from NCBI on November 29, 2019 using the update_blastdb.pl script from ncbi-blast-2.9.0+ package^19^. An initial classification of the pan-genome sequences was performed with Kaiju v1.7.2^18^. Kaiju is an amino acid-based classifier that translates nucleotide sequences and compares them to a database of proteins. As shown in the LEMMI benchmarking (https://lemmi.ezlab.org/)^36^, Kaiju have in general lower precision but good recall curves (e.g.: https://lemmi.ezlab.org/#/datasets/CAMI_I_HIGH_1/species/fig0). Because sequences are more conserved at amino acid level over large evolutionary distances, protein-based classifiers such as Kaiju allow to detect more divergent sequences. We downloaded the pre-formatted Kaiju database “NCBI BLAST *nr* +euk” that includes protein sequences from nr for Bacteria, Archaea, Viruses, Fungi and microbial eukaryotes. We additionally constructed custom Kaiju databases for other taxonomic clades corresponding to likely contaminants that are not present in the default database (Nematoda, Annelida and Platyhelminthes) and for primates. The corresponding sequences were extracted from nt/nr BLAST databases using the taxonomic information of the sequences and the program TaxonKit v0.5.0^37^. Each pan-genome was analyzed with Kaiju on each database individually using different settings: -m (minimum match length) = 11;22;33;44, and -e (number of mismatches allowed) = 3;6;9;12, respectively. The Kaiju results were then merged using the Kaiju companion script Kaiju-mergeOutputs using “-c lca” option. The outputs were converted with the companion program Kaiju2krona and visualized with krona. The results were summarized using the Kaiju2table script and full lineages of the hits were added to the tables using the Kaiju-addTaxonNames script.

Pan-genome sequences classified with Kaiju were searched in nt and nr databases with BLASTn^19^ (-evalue 1e-20) and DIAMOND v0.9.29^20^ (-evalue 1e-20) retrieving the taxonomic information of the hits (for DIAMOND using the options “--taxonmap prot.accession2taxid.gz --taxonnodes nodes.dmp”). BASTA^38^ was used to compute the LCA from the taxonomic profiles of the hits from DIAMOND (considering only those with >100 aa alignment and >70% identity from at least 3 hits) and BLASTn (>100 nucleotide alignment, >70% identity and at least 3 hits).

Prodigal v2.6.3^39^ was used to predict putative protein-coding genes of the contaminant contigs. The predicted proteins were scanned with InterProscan v5.39-77.0^40^ using the PfamA, Hamap, TIGRFAM and PANTHER databases, and additionally mapped to the bacterial node of OrthoDB v10.1^41^.

The LCAs computed with BASTA from BLASTn and DIAMOND results along with BLASTn and DIAMOND best hits and InterProscan results were merged in a table (Supplementary Table 3).

The hierarchical heat trees in figure 1 were created with the Metacoder R package^42^, using the taxonomic profiles of the contaminant sequences, i.e. the LCA computed with BASTA on DIAMOND results. The multiple sequence alignments were created and visualized with either the R package msa^43^ or the program SeaView^44^. Visualization of predicted microbial protein-coding genes on the contaminant contigs were performed with the R package Gviz^45^ by loading a track corresponding to the coordinates computed by Prodigal, and an annotation track of the orthologous group descriptions obtained from OrthoDB v10. FASTA file manipulations such as extracting, locating sequences and computing basic statistics were done using the SeqKit tool v0.11.0^46^.

## Data Availability

Supplementary tables, the list of contaminant contigs and their sequences are available in supplementary materials.

## Descriptions of Additional Supplementary Material

*Supplementary File 1*: Sequences of contaminant contigs.

*Supplementary File 2*: Predicted protein-coding genes (>40 aa) from contaminant contigs that were either mapped to an orthologous group in OrthoDB v10 or gave a hit to a Pfam domain.

*Supplementary File 3*: Graphical illustration of contaminant contigs above 10 Kbp.

*Supplementary File 4*: Soft masked sequences of the protein-coding genes identified in the Han Chinese pan-genome.

*Supplementary Table 1-2*: See Supplementary notes.

*Supplementary Table 3*: Table reporting the contaminant contigs along with their corresponding LCA classifications and best hits from DIAMOND/BLASTn, and annotations from Pfam and Interpro databases.

*Supplementary Table 4*: Interproscan results of the predicted proteins (> 40 aa) from contaminant contigs.

*Supplementary Table 5*: Accessions and publication dates of microbial sequences deposited in NCBI that matched the contaminant contigs. Only the three oldest sequences are reported. This data was used for the plot of figure 1 Panel D.

*Supplementary Table 6*: Table reporting the percentage of masked sequence for each of the 188 protein-coding genes from the Han Chinese pan-genome.

*Supplementary Table 7*: Statistic of RepeatMasker on the 188 protein-coding genes from the Han Chinese pan-genome.

*Supplementary Table 8*: Coordinates of repetitive regions masked by the program dna-brnn on the 188 protein-coding genes from the Han Chinese pan-genome.

*Supplementary Table 9*: LCA and best hits of the seven protein-coding genes from the Han Chinese pan-genome with similarity to bacterial proteins.

*Supplementary Table 10*: List of bona fide human sequences that match hypothetical proteins of Plasmodium ovale wallikeri.

## Acknowledgments

We thank Fredrik T for mapping data to OrthoDB orthologous groups. Grazia S and Chris R for useful comments on the work.

## Author Contributions

MM and EZM managed and coordinated the study; MM wrote the manuscript, prepared figures and tables. All authors read, corrected and commented on the manuscript.

## Competing Interests

The authors declare no competing financial interests.

