## Supplementary_materials for "Microbial contaminants cataloged as novel human sequences in recent human pan-genomes": Supplementary_File_3.pdf

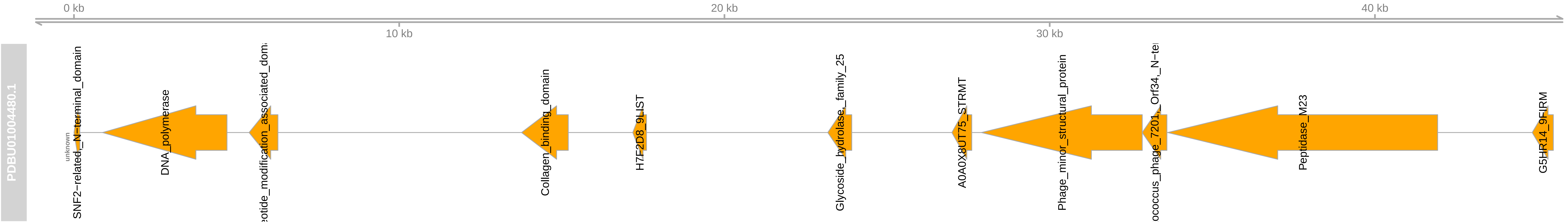

0 kb
10 kb
20 kb
30 kb
40 kb
PDBU01004480.1
unknown
SNF2−related_N−terminal_domain
DNA_polymerase
Collagen_binding_domain
H7F2D8_9LIST
Glycoside_hydrolase_family_25
A0A0X8UT75_STRMT
Phage_minor_structural_protein
Peptidase_M23
G5HR14_9FIRM

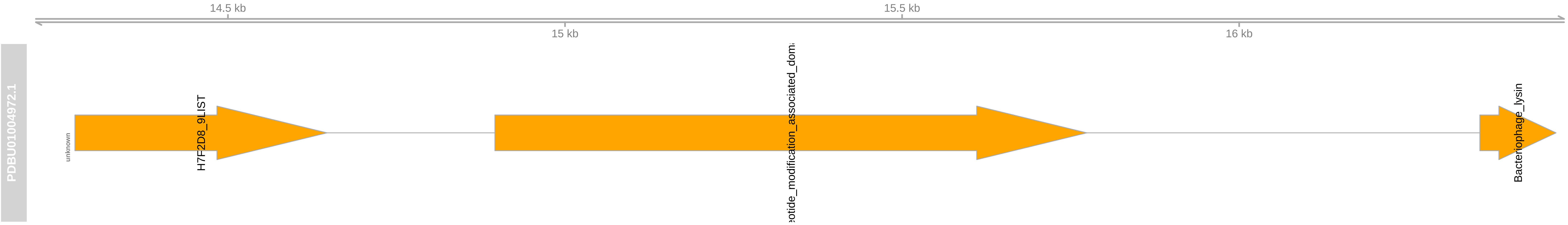
14.5 kb
15.5 kb
15 kb
16 kb
PDBU01004972.1
unknown
H7F2D8_9LIST
eotide_modification_associated_dom
Bacteriophage_lysin

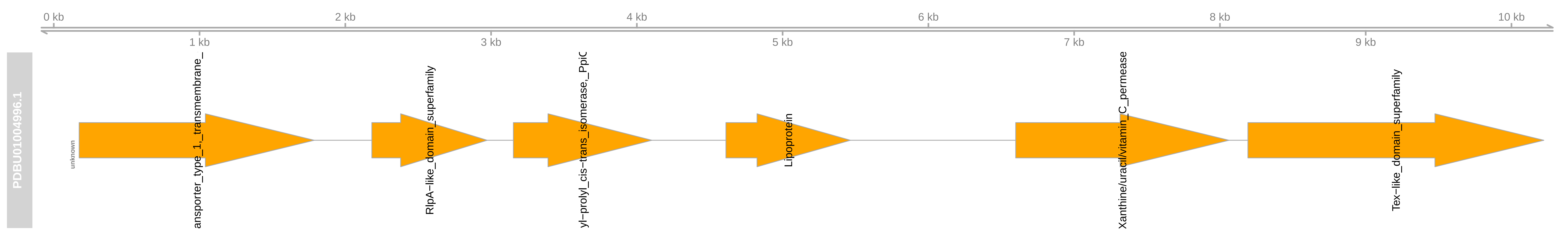
0 kb
1 kb
2 kb
3 kb
4 kb
5 kb
6 kb
7 kb
8 kb
9 kb
10 kb
PDBU01004996.1
unknown
ansporter_type_1,_transmembrane_
RlpA−like_domain_superfamily
yl−prolyl_cis−trans_isomerase,_PpiC
Lipoprotein
Xanthine/uracil/vitamin_C_permease
Tex−like_domain_superfamily

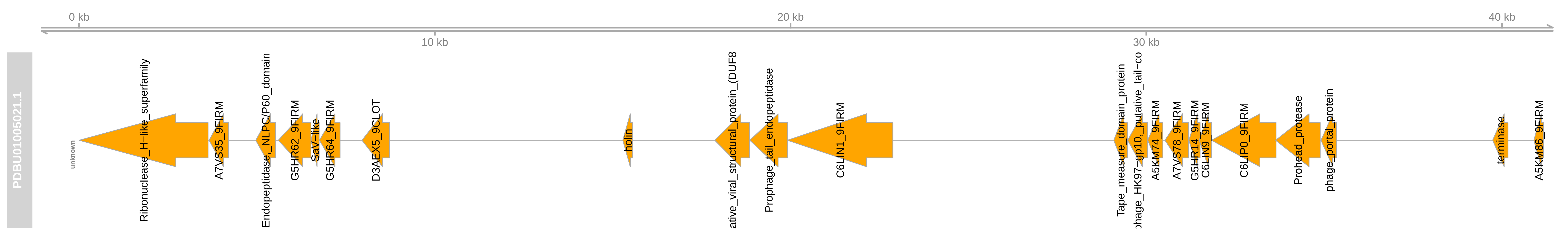

0 kb
10 kb
20 kb
30 kb
40 kb
PDBU01005021.1
unknown
Ribonuclease_H-like_superfamily
A7VS35_9FIRM
Endopeptidase_NLPC/P60_domain
G5HR62_9FIRM
SaV-like
G5HR64_9FIRM
D3AEX5_9CLOT
holin
ative_viral_structural_protein_(DUF8
Prophage_tail_endopeptidase
C6LIN1_9FIRM
Tape_measure_domain_protein
phage_HK97-gp10_putative_tail-co
A5KM74_9FIRM
A7VS78_9FIRM
G5HR14_9FIRM
C6LIN9_9FIRM
C6LIP0_9FIRM
Prohead_protease
phage_portal_protein
terminase
A5KM86_9FIRM

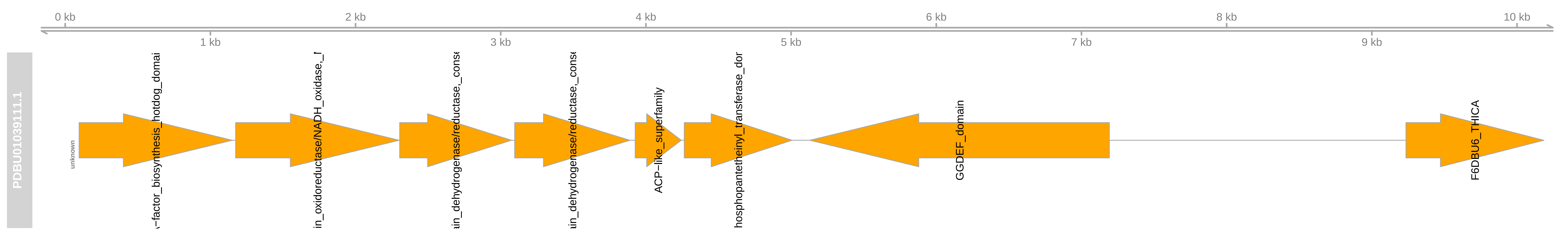
0 kb
1 kb
2 kb
3 kb
4 kb
5 kb
6 kb
7 kb
8 kb
9 kb
10 kb
PDBU01039111.1
unknown
–factor_biosynthesis_hotdog_domai
in_oxidoreductase/NADH_oxidase,_f
ain_dehydrogenase/reductase,_cons
ain_dehydrogenase/reductase,_cons
ACP–like_superfamily
hosphopantetheinyl_transferase_dor
GGDEF_domain
F6DBU6_THICA

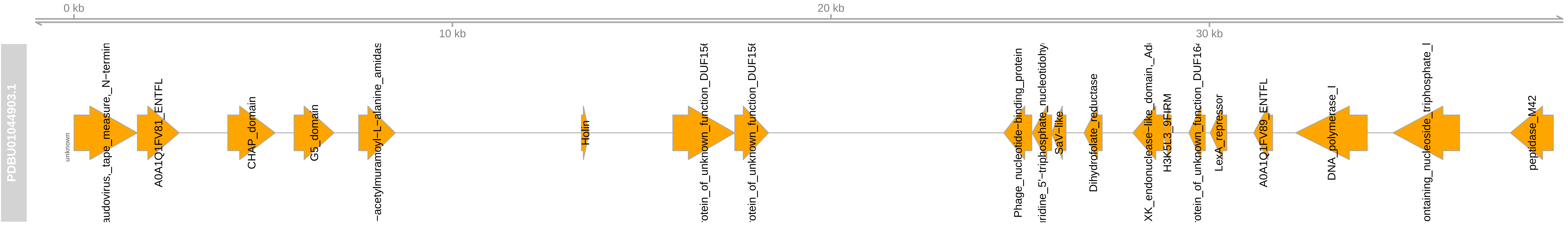
0 kb
10 kb
20 kb
30 kb
PDBU01044903.1
unknown
audovirus,_tape_measure,_N-termin
A0A1Q1FV81_ENTFL
CHAP_domain
G5_domain
-acetylmuramoyl-L-alanine_amidas
Holin
rotein_of_unknown_function_DUF15
rotein_of_unknown_function_DUF15
Phage_nucleotide-binding_protein
ridine_5'-triphosphate_nucleotidohy
SaV-like
Dihydrofolate_reductase
XK_endonuclease-like_domain,_Ad
H3K5L3_9FIRM
rotein_of_unknown_function_DUF16
LexA_repressor
A0A1Q1FV89_ENTFL
DNA_polymerase_I
ontaining_nucleoside_triphosphate_I
peptidase_M42

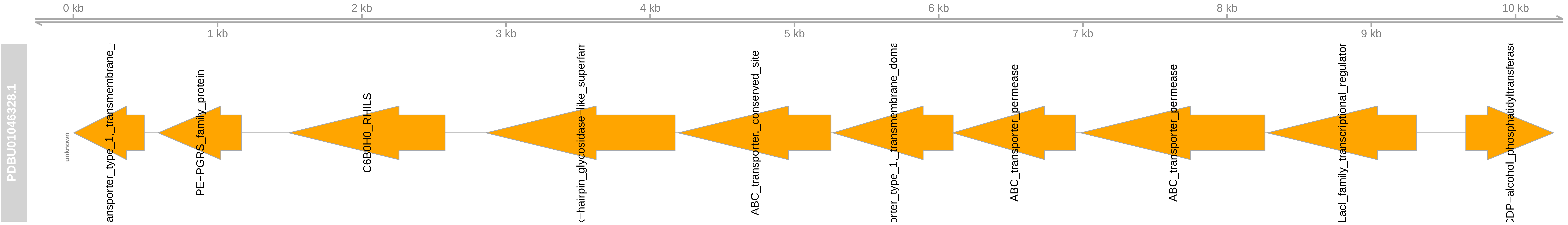
0 kb
2 kb
4 kb
6 kb
8 kb
10 kb
1 kb
3 kb
5 kb
7 kb
9 kb
PDBU01046328.1
unknown
ansporter_type_1,_transmembrane_
PE−PGRS_family_protein
C6B0H0_RHILS
−hairpin_glycosidase−like_superfam
ABC_transporter,_conserved_site
orter_type_1,_transmembrane_dom
ABC_transporter_permease
ABC_transporter_permease
LacI_family_transcriptional_regulator
CDP−alcohol_phosphatidyltransferase

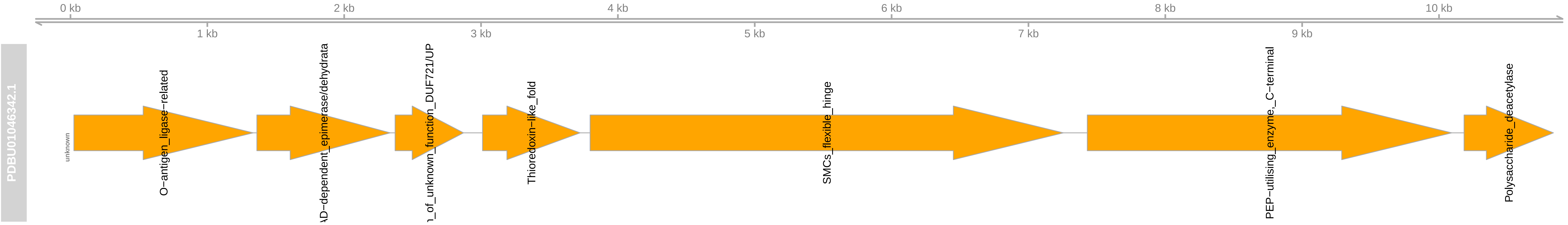
0 kb
2 kb
4 kb
6 kb
8 kb
10 kb
1 kb
3 kb
5 kb
7 kb
9 kb
PDBU01046342.1
unknown
O−antigen_ligase−related
AD−dependent_epimerase/dehydrata
n_of_unknown_function_DUF721/UP
Thioredoxin−like_fold
SMCs_flexible_hinge
PEP−utilising_enzyme,_C−terminal
Polysaccharide_deacetylase

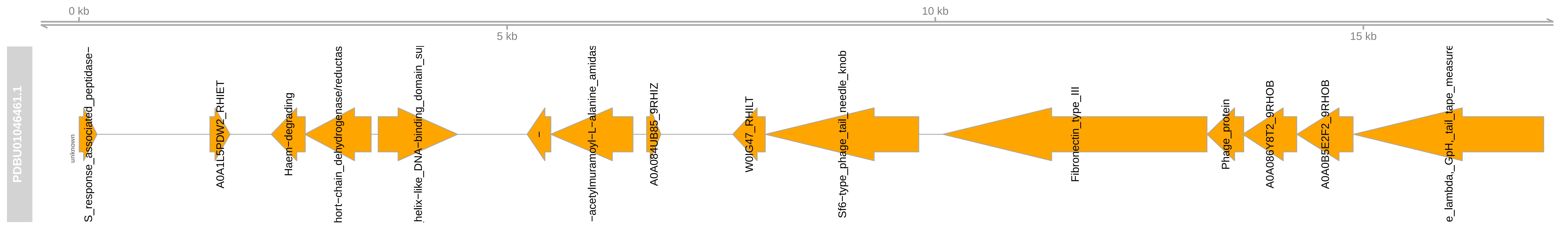
0 kb
5 kb
10 kb
15 kb
PDBU01046461.1
unknown
S_response_associated_peptidase–
A0A1L5PDW2_RHIET
Haem–degrading
hort–chain_dehydrogenase/reductas
helix–like_DNA–binding_domain_su
–
–acetylmuramoyl–L–alanine_amidas
A0A084UB85_9RHIZ
W0IG47_RHILT
Sf6–type_phage_tail_needle_knob
Fibronectin_type_III
Phage_protein
A0A086Y8T2_9RHOB
A0A0B5E2F2_9RHOB
e_lambda,_GpH,_tail_tape_measure

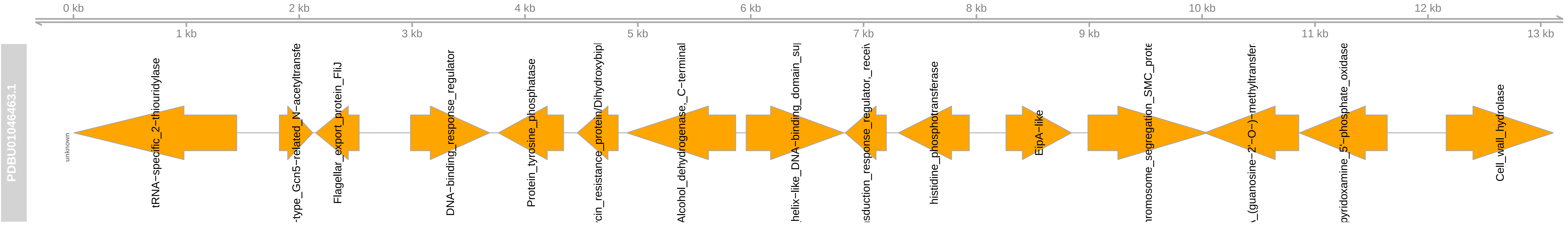

0 kb
2 kb
4 kb
6 kb
8 kb
10 kb
12 kb
1 kb
3 kb
5 kb
7 kb
9 kb
11 kb
13 kb
PDBU01046463.1
unknown
tRNA−specific_2−thiouridylase
Flagellar_export_protein_FliJ
DNA−binding_response_regulator
Protein_tyrosine_phosphatase
Alcohol_dehydrogenase,_C−terminal
histidine_phosphotransferase
EipA−like
Cell_wall_hydrolase

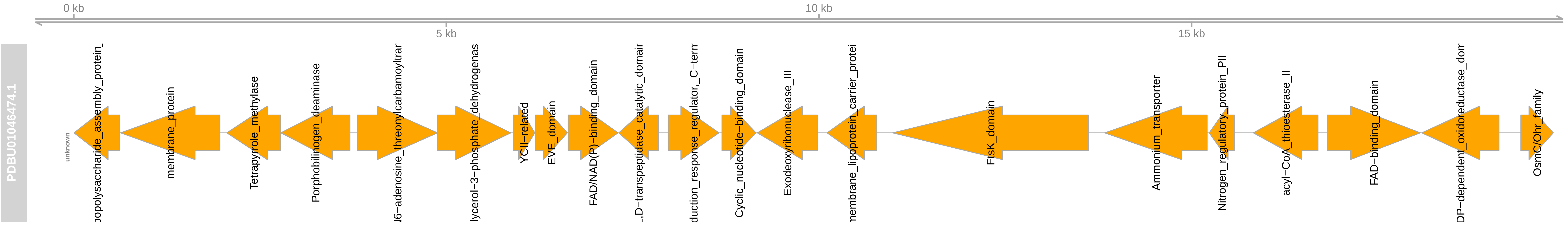

0 kb
5 kb
10 kb
15 kb
PDBU01046474.1
unknown
opolysaccharide_assembly_protein_
membrane_protein
Tetrapyrrole_methylase
Porphobilinogen_deaminase
N6−adenosine_threonylcarbamoyltrar
lycerol−3−phosphate_dehydrogenas
YCII−related
EVE_domain
FAD/NAD(P)−binding_domain
..D−transpeptidase_catalytic_domair
duction_response_regulator,_C−term
Cyclic_nucleotide−binding_domain
Exodeoxyribonuclease_III
nembrane_lipoprotein_carrier_protei
FtsK_domain
Ammonium_transporter
Nitrogen_regulatory_protein_PII
acyl−CoA_thioesterase_II
FAD−binding_domain
DP−dependent_oxidoreductase_dom
OsmC/Ohr_family

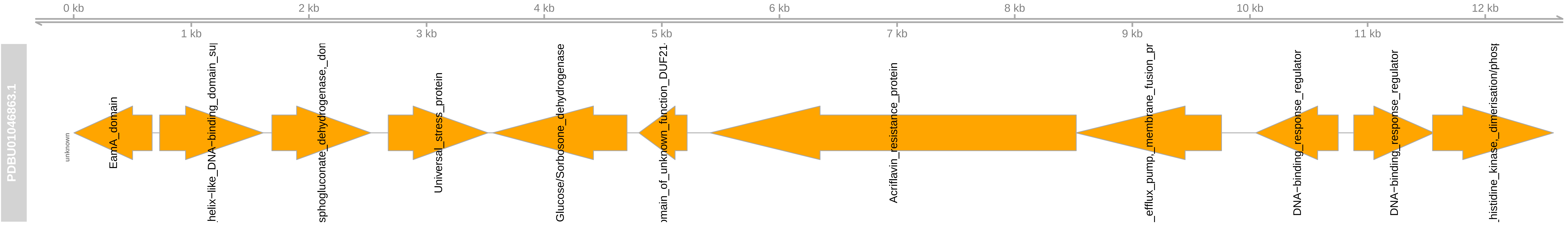
0 kb
1 kb
2 kb
3 kb
4 kb
5 kb
6 kb
7 kb
8 kb
9 kb
10 kb
11 kb
12 kb
PDBU01046863.1
unknown
EamA_domain
helix-like_DNA-binding_domain_su
sphogluconate_dehydrogenase,_dom
Universal_stress_protein
Glucose/Sorbosone_dehydrogenase
omain_of_unknown_function_DUF21
Acriflavin_resistance_protein
_efflux_pump,_membrane_fusion_pr
DNA-binding_response_regulator
DNA-binding_response_regulator
_histidine_kinase,_dimerisation/phos

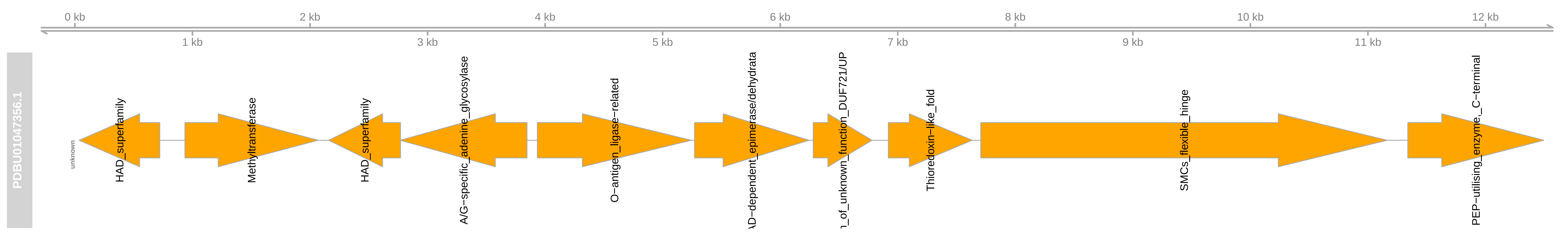
0 kb
2 kb
4 kb
6 kb
8 kb
10 kb
12 kb
1 kb
3 kb
5 kb
7 kb
9 kb
11 kb
PDBU01047356.1
unknown
HAD_superfamily
Methyltransferase
HAD_superfamily
A/G−specific_adenine_glycosylase
O−antigen_ligase−related
AD−dependent_epimerase/dehydrata
n_of_unknown_function_DUF721/UP
Thioredoxin−like_fold
SMCs_flexible_hinge
PEP−utilising_enzyme,_C−terminal

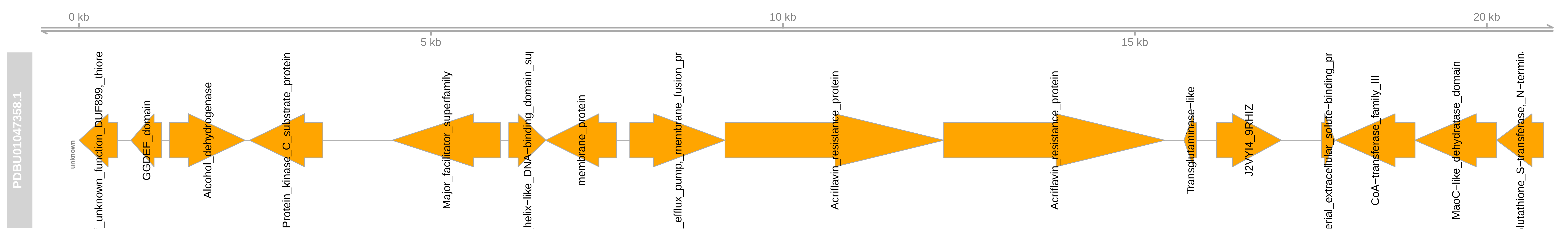
0 kb
5 kb
10 kb
15 kb
20 kb
PDBU01047358.1
unknown
_unknown_function_DUF899,_thiore
GGDEF_domain
Alcohol_dehydrogenase
Protein_kinase_C_substrate_protein
Major_facilitator_superfamily
helix−like_DNA−binding_domain_su
membrane_protein
efflux_pump,_membrane_fusion_pr
Acriflavin_resistance_protein
Acriflavin_resistance_protein
Transglutaminase−like
J2VYI4_9RHIZ
erial_extracellular_solute−binding_pr
CoA−transferase_family_III
MaoC−like_dehydratase_domain
lutathione_S−transferase,_N−termin

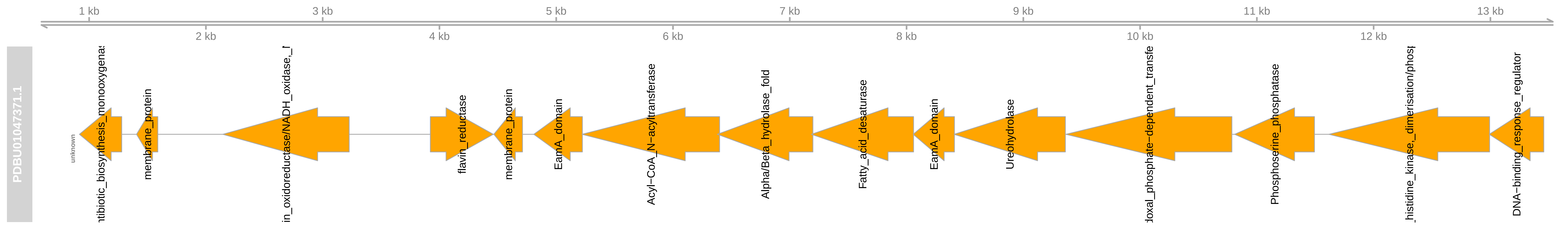
1 kb
3 kb
5 kb
7 kb
9 kb
11 kb
13 kb
2 kb
4 kb
6 kb
8 kb
10 kb
12 kb
PDBU01047371.1
unknown
ntibiotic_biosynthesis_monooxygenas
membrane_protein
in_oxidoreductase/NADH_oxidase,_f
flavin_reductase
membrane_protein
EamA_domain
Acyl−CoA_N−acyltransferase
Alpha/Beta_hydrolase_fold
Fatty_acid_desaturase
EamA_domain
Ureohydrolase
doxal_phosphate−dependent_transfe
Phosphoserine_phosphatase
_histidine_kinase,_dimerisation/phosp
DNA−binding_response_regulator

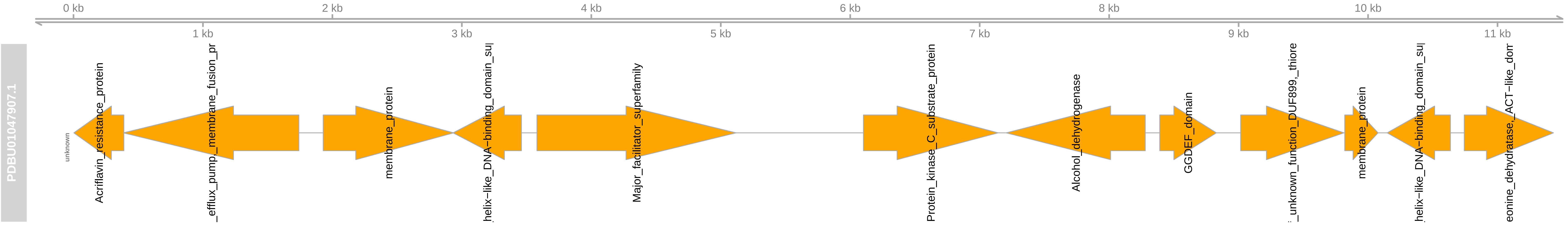

0 kb
2 kb
4 kb
6 kb
8 kb
10 kb
1 kb
3 kb
5 kb
7 kb
9 kb
11 kb
PDBU01047907.1
unknown
Acriflavin_resistance_protein
_efflux_pump_membrane_fusion_pr
membrane_protein
helix-like_DNA-binding_domain_su
Major_facilitator_superfamily
Protein_kinase_C_substrate_protein
Alcohol_dehydrogenase
GGDEF_domain
_unknown_function_DUF899,_thiore
membrane_protein
helix-like_DNA-binding_domain_su
eonine_dehydratase,_ACT-like_dom

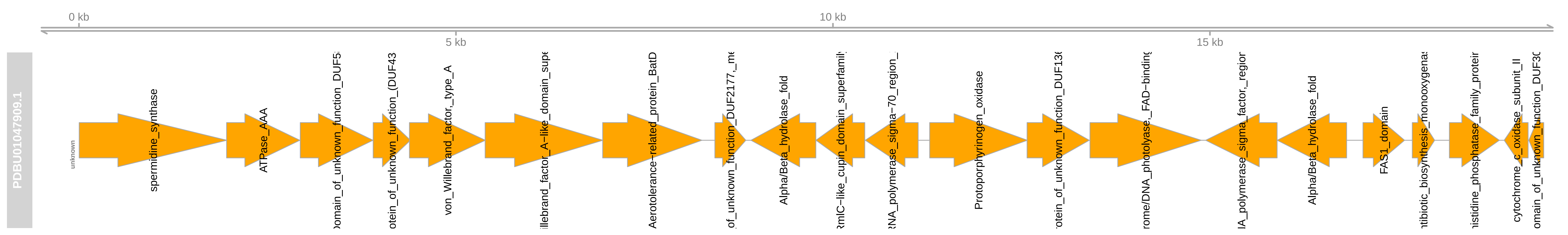
0 kb
5 kb
10 kb
15 kb
PDBU01047909.1
unknown
spermidine_synthase
ATPase_AAA
omain_of_unknown_function_DUF5
otein_of_unknown_function_(DUF43
von_Willebrand_factor,_type_A
illebrand_factor_A-like_domain_supe
Aerotolerance-related_protein_BatD
of_unknown_function_DUF2177,_me
Alpha/Beta_hydrolase_fold
RmlC-like_cupin_domain_superfamily
RNA_polymerase_sigma-70_region_
Protoporphyrinogen_oxidase
otein_of_unknown_function_DUF136
rome/DNA_photolyase,_FAD-binding
A_polymerase_sigma_factor,_regior
Alpha/Beta_hydrolase_fold
FAS1_domain
tibiotic_biosynthesis_monooxygena
istidine_phosphatase_family_protei
cytochrome_c_oxidase_subunit_II
omain_of_unknown_function_DUF30

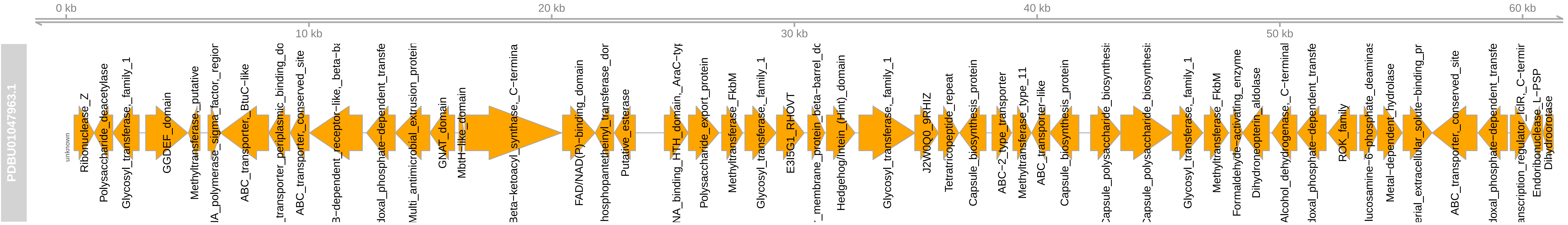

0 kb
10 kb
20 kb
30 kb
40 kb
50 kb
60 kb
PDBU01047963.1
unknown
Ribonuclease_Z
Polysaccharide_deacetylase
Glycosyl_transferase,_family_1
GGDEF_domain
Methyltransferase,_putative
ABC_transporter,_BtuC-like
ABC_transporter,_conserved_site
GNAT_domain
MbtH-like_domain
FAD/NAD(P)-binding_domain
Putative_esterase
Polysaccharide_export_protein
Methyltransferase_FkbM
Glycosyl_transferase,_family_1
E3I5G1_RHOVT
Hedgehog/intein_(Hint)_domain
Glycosyl_transferase,_family_1
J2W0Q0_9RHIZ
Tetratricopeptide_repeat
Capsule_biosynthesis_protein
ABC-2_type_transporter
Methyltransferase_type_11
ABC_transporter-like
Capsule_biosynthesis_protein
Glycosyl_transferase,_family_1
Methyltransferase_FkbM
Formaldehyde-activating_enzyme
Dihydroneopterin_aldolase
ROK_family
Metal-dependent_hydrolase
ABC_transporter,_conserved_site
Endoribonuclease_L-PSP
Dihydroorotase

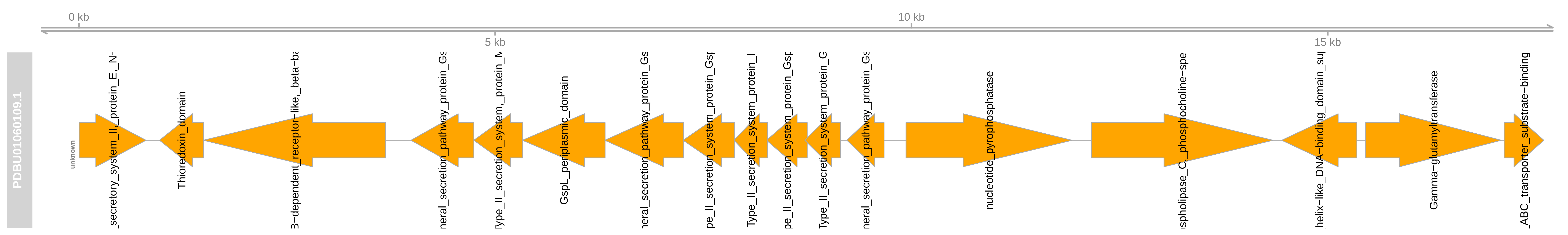
0 kb
5 kb
10 kb
15 kb
PDBU01060109.1
unknown
_secretory_system_II,_protein_E,_N-
Thioredoxin_domain
3-dependent_receptor-like,_beta-ba
neral_secretion_pathway_protein_Gs
Type_II_secretion_system,_protein_M
GspL_periplasmic_domain
neral_secretion_pathway_protein_Gs
pe_II_secretion_system_protein_Gsp
Type_II_secretion_system_protein_I
pe_II_secretion_system_protein_Gsp
Type_II_secretion_system_protein_G
neral_secretion_pathway_protein_Gs
nucleotide_pyrophosphatase
spholipase_C,_phosphocholine-spe
helix-like_DNA-binding_domain_su
Gamma-glutamyltransferase
_ABC_transporter_substrate-binding

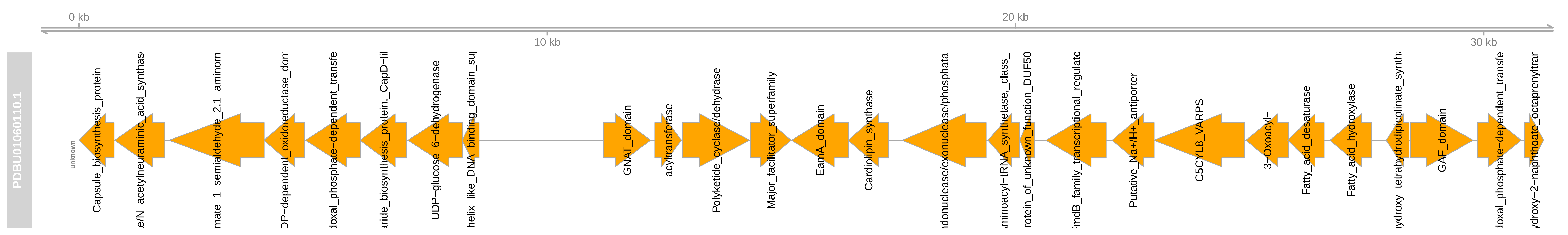
0 kb
10 kb
20 kb
30 kb
PDBU01060110.1
unknown
Capsule_biosynthesis_protein
e/N−acetylneuraminic_acid_synthas
mate−1−semialdehyde_2,1−aminom
DP−dependent_oxidoreductase_dom
doxal_phosphate−dependent_transfe
aride_biosynthesis_protein,_CapD−li
UDP−glucose_6−dehydrogenase
helix−like_DNA−binding_domain_su
GNAT_domain
acyltransferase
Polyketide_cyclase/dehydrase
Major_facilitator_superfamily
EamA_domain
Cardiolipin_synthase
ndonuclease/exonuclease/phosphata
minoacyl−tRNA_synthetase,_class_
rotein_of_unknown_function_DUF50
rmB_family_transcriptional_regulato
Putative_Na+/H+_antiporter
C5CYL8_VARPS
3−Oxoacyl−
Fatty_acid_desaturase
Fatty_acid_hydroxylase
ydroxy−tetrahydrodipicolinate_synth
GAF_domain
doxal_phosphate−dependent_transfe
ydroxy−2−naphthoate_octaprenyltrar

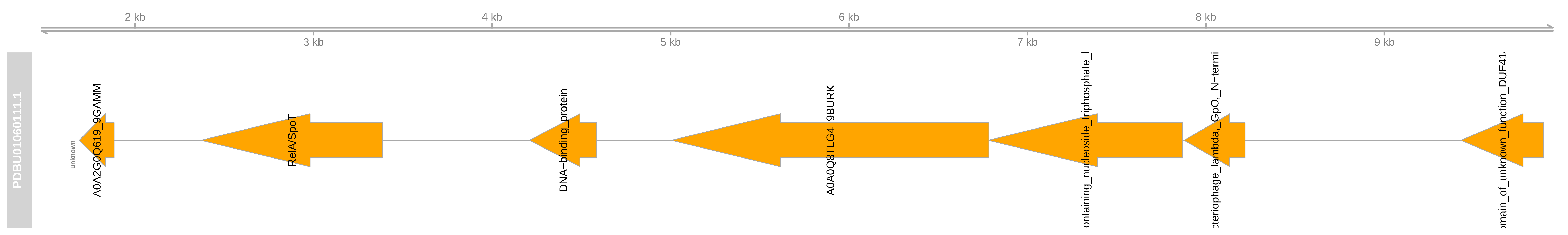

2 kb
4 kb
6 kb
8 kb
3 kb
5 kb
7 kb
9 kb
PDBU01060111.1
unknown
A0A2G0Q619_9GAMM
RelA/SpoT
DNA−binding_protein
A0A0Q8TLG4_9BURK
ontaining_nucleoside_triphosphate_l
cteriophage_lambda,_GpO,_N−termi
omain_of_unknown_function_DUF41.

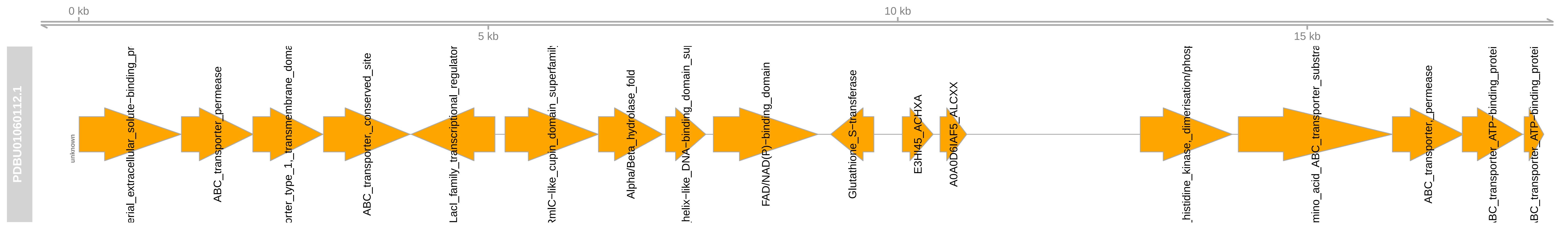

0 kb
10 kb
5 kb
15 kb
PDBU01060112.1
unknown
erial_extracellular_solute−binding_pr
ABC_transporter_permease
orter_type_1,_transmembrane_doma
ABC_transporter,_conserved_site
LacI_family_transcriptional_regulator
RmlC−like_cupin_domain_superfamily
Alpha/Beta_hydrolase_fold
helix−like_DNA−binding_domain_sup
FAD/NAD(P)−binding_domain
Glutathione_S−transferase
E3HI45_ACHXA
A0A0D6IAF5_ALCXX
histidine_kinase,_dimerisation/phosp
mino_acid_ABC_transporter_substra
ABC_transporter,_permease
ABC_transporter_ATP−binding_protei
ABC_transporter_ATP−binding_protei

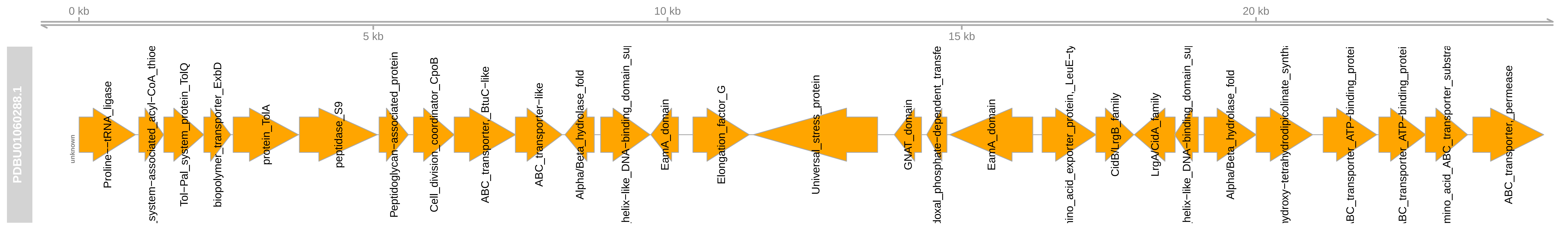

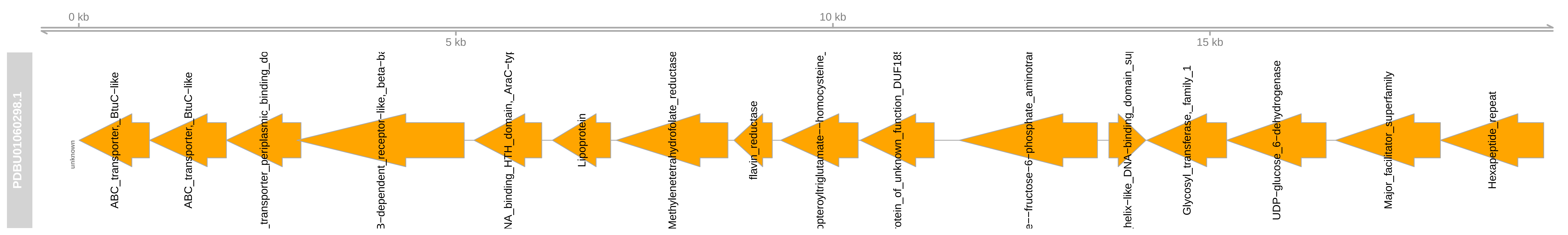
0 kb
10 kb
5 kb
15 kb
PDBU01060298.1
unknown
ABC_transporter,_BtuC-like
ABC_transporter,_BtuC-like
transporter_periplasmic_binding_do
B-dependent_receptor-like,_beta-ba
NA_binding_HTH_domain,_AraC-typ
Lipoprotein
Methylenetetrahydrofolate_reductase
flavin_reductase
opteroyltriglutamate--homocysteine_
otein_of_unknown_function_DUF18
e--fructose-6-phosphate_aminotra
helix-like_DNA-binding_domain_su
Glycosyl_transferase,_family_1
UDP-glucose_6-dehydrogenase
Major_facilitator_superfamily
Hexapeptide_repeat

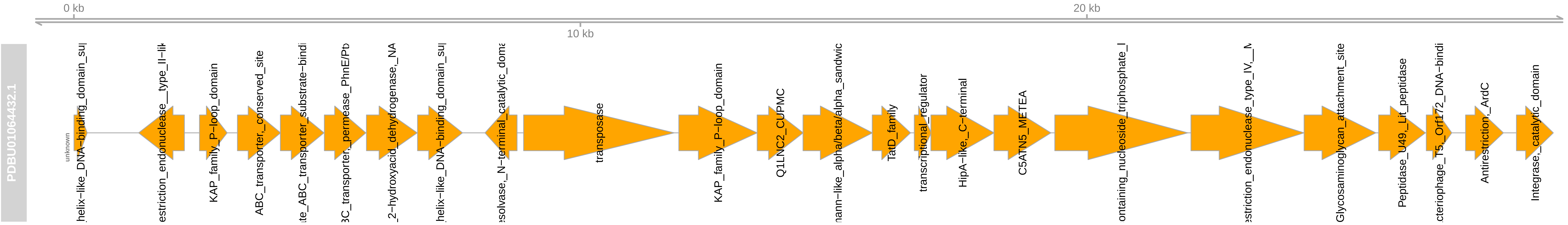
0 kb
10 kb
20 kb
PDBU01064432.1
unknown
helix−like_DNA−binding_domain_su
estriction_endonuclease__type_II−lik
KAP_family_P−loop_domain
ABC_transporter,_conserved_site
ate_ABC_transporter_substrate−bindi
BC_transporter,_permease_PhnE/Pt
2−hydroxyacid_dehydrogenase,_NA
helix−like_DNA−binding_domain_su
esolvase,_N−terminal_catalytic_doma
transposase
KAP_family_P−loop_domain
Q1LNC2_CUPMC
nann−like_alpha/beta/alpha_sandwic
TatD_family
transcriptional_regulator
HipA−like,_C−terminal
C5ATN5_METEA
ontaining_nucleoside_triphosphate_l
estriction_endonuclease_type_IV,__M
Glycosaminoglycan_attachment_site
Peptidase_U49,_Lit_peptidase
cteriophage_T5,_Orf172_DNA−bindi
Antirestriction,_ArdC
Integrase,_catalytic_domain

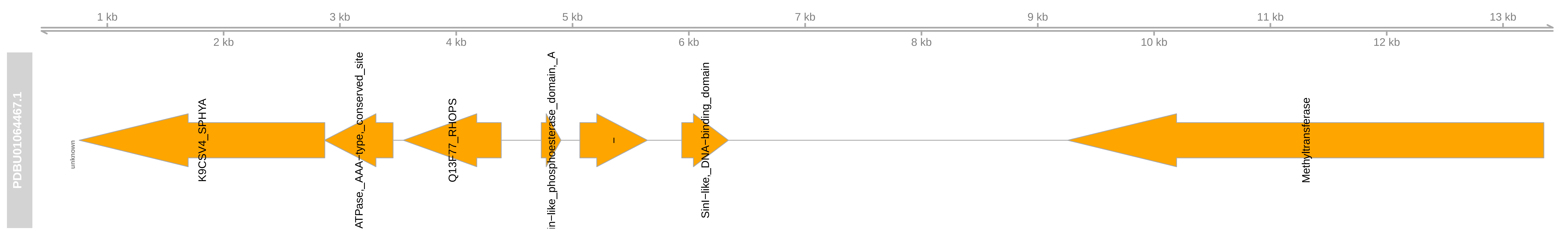

1 kb
3 kb
5 kb
7 kb
9 kb
11 kb
13 kb
2 kb
4 kb
6 kb
8 kb
10 kb
12 kb
PDBU01064467.1
unknown
K9CSV4_SPHYA
ATPase,_AAA−type,_conserved_site
Q13F77_RHOPS
in−like_phosphoesterase_domain,_A
−
SinI−like,_DNA−binding_domain
Methyltransferase

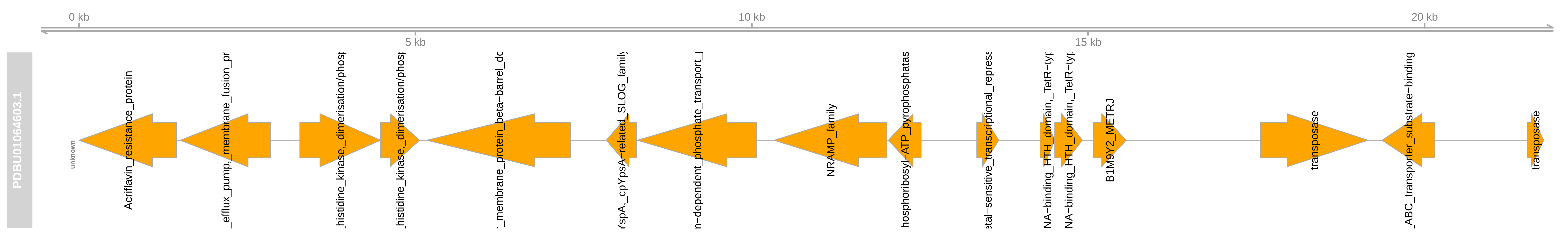
0 kb
5 kb
10 kb
15 kb
20 kb
PDBU01064603.1
unknown
Acriflavin_resistance_protein
_efflux_pump,_membrane_fusion_pr
histidine_kinase,_dimerisation/phosp
histidine_kinase,_dimerisation/phosp
membrane_protein_beta-barrel_dc
YspA,_cpYpsA-related_SLOG_family
n-dependent_phosphate_transport_
NRAMP_family
hosphoribosyl-ATP_pyrophosphatas
etal-sensitive_transcriptional_repress
NA-binding_HTH_domain,_TetR-typ
NA-binding_HTH_domain,_TetR-typ
B1M9Y2_METRJ
transposase
_ABC_transporter_substrate-binding
transposase

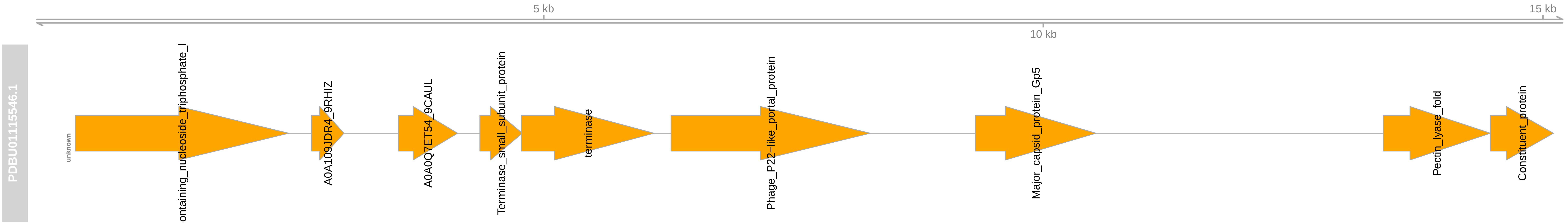
5 kb
10 kb
15 kb
PDBU01115546.1
unknown
ontaining_nucleoside_triphosphate_l
A0A109JDR4_9RHIZ
A0A0Q7ET54_9CAUL
Terminase_small_subunit_protein
terminase
Phage_P22-like_portal_protein
Major_capsid_protein_Gp5
Pectin_lyase_fold
Constituent_protein

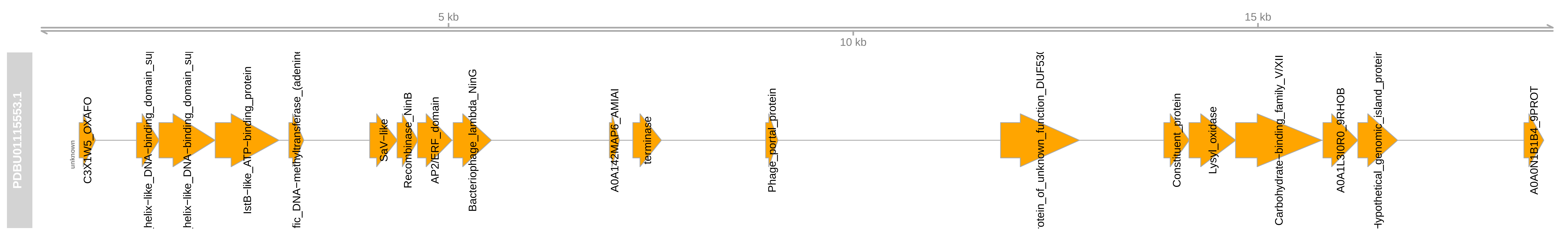
5 kb
10 kb
15 kb
PDBU01115553.1
unknown
C3X1W5_OXAFO
helix−like_DNA−binding_domain_su
helix−like_DNA−binding_domain_su
IstB−like_ATP−binding_protein
fic_DNA−methyltransferase_(adenine
SaV−like
Recombinase_NinB
AP2/ERF_domain
Bacteriophage_lambda_NinG
A0A142MAP6_AMIAI
terminase
Phage_portal_protein
rotein_of_unknown_function_DUF53
Constituent_protein
Lysyl_oxidase
Carbohydrate−binding_family_V/XII
A0A1L3I0R0_9RHOB
Hypothetical_genomic_island_protei
A0A0N1B1B4_9PROT
