## Supplementary_materials for "Microbial contaminants cataloged as novel human sequences in recent human pan-genomes": Supplementary_notes.pdf

The African pan-genome was constructed from a dataset of 910 deeply sequenced individuals[1]. The pan-genome consist of 296 Mb in ~126,000 contigs. The authors were able to anchor 1,548 contigs (4.4 Mbp in total) to a specific location in the GRCh38 assembly, and reported that 387 contigs fall within 315 protein-coding genes. As example of contaminant contigs, at least 70 contigs gave hit to *Burkholderia stabilis* with a >99% nucleotide identity over >1,000 bp alignment, and several other contigs to *Burkholderia contaminans*. The majority of these sequences harbour protein coding genes. As example, a 887 aa long predicted protein from contig PDBU01039112.1 matches *Burkholderia stabilis* fimbrial biogenesis outer membrane usher protein (WP_156814889.1, 100% identity over the all sequence), which is involved in biogenesis of the pilus in Gram-negative bacteria. Around 250 contigs are related to the order Rhyzobiales among which *Bradyrhizobium* and *Methylobacteria* genuses. PDBU01069001.1 (2,063 bp) aligns with a sequences of *Bradyrhizobium sp. SK17*. The contig encodes for a partial protein of 687 aa (PDBU01069001.1_1) which has 100% identity to a MMPL family transporter of *Bradyrhizobium* sp. MOS004. At least 50 contigs share between 90-100% similarity over >1,000 bp with *Mogibacterium diversum* a Gram-positive bacterium which has been isolated from the human mouth[2]. More than 100 contigs resemble a *Gemella* species, with best hit to *Gemella haemolysans*. For the majority of the sequences, though the nucleotide identity of the alignments ranged from 70-99%.

The Han Chinese pan-genome was compiled from 275 genomes, including 185 newly sequenced[3] and 90 previously published genomes[4]. Overall the study identified 29.50 Mb of high-confidence sequences missing from the GRCh38 (including patch sequences and alternative loci) and 188 novel human genes. By comparing these sequences to other pan-genome resources, including the APG, the authors identified 13.26 Mb of sequences novel and specific to Han Chinese individuals. Almost all contaminants gave hit to sequences of *Kocuria indica* with >95% nucleotide similarity. *Kocuria indica* is a high-GC-content Gram-positive aerobic bacterium belonging to the family *Micrococcaceae* in the phylum Actinobacteria that have been isolated from environmental samples[5] and human skin[6]. The contaminant contigs are more closely related to the latter. This assembly (GCF_004328535.1) was sequenced from cultured cells using PacBio RSII long reads at 124-fold genome coverage. The assembly is of high quality and unlikely to contain human sequence contaminants. The contaminant contigs also match sequences from other *Kocuria* species and the LCA analysis strongly suggests a species belonging to *Kokuria* is the likely origin of these contigs. As example of contaminant contig, HCP020618 share a 99% similarity over 1,742 base pair with *Kocuria indica* strain CE7 (CP035504.1), while the second best hit is with *Kocuria sp*. BT304 with a lower nucleotide identity of 75%. HCP020618 encodes a predicted protein (HCP020618_2; 262 aa) of the type 2 secretion system, a secretion machinery found in various species of Gram-negative bacteria.

**Table S1:** Source of pan-genome data used in the analyses.

| Pangenome | Abbreviation | Version | Download date | Source |
|---|---|---|---|---|
| African pan-genome | APG | PDBU01000000 | 15 Dec. 2019 | https://www.ncbi.nlm.nih.gov/assembly/GCA_003610035.1/ |
| Han Chinese pan-genome | HCP | - | 15 Dec. 2019 | http://cgm.sjtu.edu.cn/hupan/src/HanChinesePan.fa.gz<br>http://cgm.sjtu.edu.cn/hupan/data/combined.188genes.minus_patch_and_alternative.gene.fasta |
| Swedish NRSs | Swed NRSs | 20180409 | 15 Dec. 2019 | https://swefreq.nbis.se<br>https://swefreq.nbis.se/release/SweGen/versions/20180409/9eaecf954eee4bf4ba3879697ed7c1b5/swegen_novel_sequences.zip |

**Table S2:** Statistics of masked sequences.

| Sequence | Size (bp) | Total size masked (bp) [% size] | Size masked by Repeat Masker (bp) [% size] |
|---|---|---|---|
| APG | 296,485,284 | 283,908,766 [95] | 268,903,065 [90.7] |
| HCP | 29,503,170 | 25,957,964 [87] | 23,062,689 [78.2] |
| Swed NRSs | 46,839,038 | 30,854,530 [65] | 25,524,874 [54.5] |

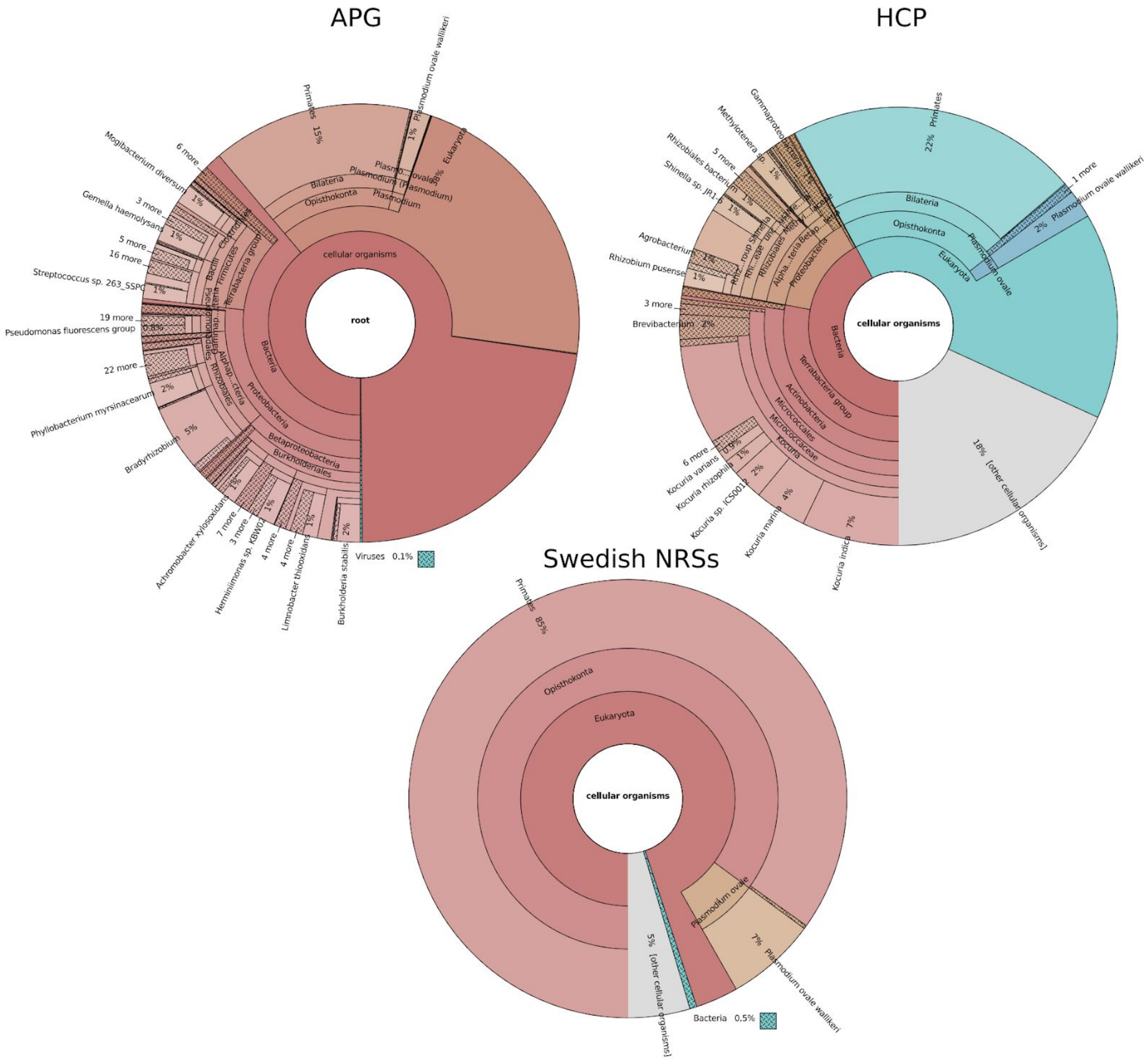


**Supplementary Figure 1.** Visualization of the initial results from Kaiju.
